# Transient inhibition of cell division in competent pneumococcal cells results from deceleration of the septal peptidoglycan complex

**DOI:** 10.1101/2024.02.28.582556

**Authors:** Dimitri Juillot, Cyrille Billaudeau, Aurélien Barbotin, Armand Lablaine, Isabelle Mortier-Barrière, Patrice Polard, Nathalie Campo, Rut Carballido-Lopez

## Abstract

Bacterial cells are known to produce inhibitors of cell division in response to stress responses and developmental programs. Knowledge of the underlying molecular mechanisms remains however largely limited. In this study, we investigated the mechanism of transient cell division inhibition observed during the development of competence for transformation in the human pathogen *Streptococcus pneumoniae*. In this species, ComM, a membrane protein specifically produced during competence, transiently inhibits cell division to preserve genomic integrity during transformation. We show that ComM reduces specifically the dynamics of the septal peptidoglycan synthetic complex FtsW:PBP2x. We also present evidence that ComM interacts with the peptidoglycan precursor synthetic enzyme MurA, and show that overproduction of MurA suppresses FtsW:PBP2x deceleration along the cell division delay in competent cells. Collectively, our data support a model in which ComM interferes with MurA activity to reduce septal peptidoglycan synthesis during competence in *S. pneumoniae*.

---

In bacteria, environmental and cellular stresses including nutrient exhaustion, envelope stress and DNA damages, trigger various signaling pathways that can cause a transient inhibition of the cell division process to halt the cell cycle progression and allow time for repair^1,2^. Inhibition of cell division is also frequently associated to differentiation programs such as sporulation in *Bacillus subtilis*^3^, heterocysts formation in Anabaena^4,5^ and aerial hyphal development in Streptomyces^6–8^. However, most cellular factors and mechanisms arresting cell division in response to specific conditions that enable bacteria to adapt remain largely unknown. The best characterized mechanism involves the SOS cell division inhibitor SulA in *Escherichia coli*, which is induced following activation of the RecA protein by damage to DNA^9^. SulA binds to monomers of the bacterial tubulin homolog FtsZ in the cytoplasm, and prevents their polymerization into a ring-like structure (Z-ring) in the membrane^10^. The Z-ring marks the site of cell division and directs the assembly of the cell division machinery^11^. It appears however that, while other factors such as MciZ in *B. subtilis*^12^ also impact directly the polymerization of FtsZ, most of the known cell division inhibitors localize in the membrane and interfere with the activities of various components of the cell division machinery^13–19^.

This machinery, also called the divisome^11,20^, is a large multiprotein complex containing proteins involved in cell division and cell wall peptidoglycan (PG) synthesis that collectively achieve membrane invagination during cytokinesis and build a crosswall (the division septum) that will be split to give rise to two daughter cells. Among the divisome components, FtsZ plays a major role by coordinating the cell constriction process. Several studies have demonstrated that FtsZ filaments treadmill circumferentially around the Z-ring with an average velocity of ∼30 nm/s ^21–25^. In the rod-shaped *B. subtilis* and *E. coli* bacteria, the dynamics of FtsZ filaments drives the motions of PG-synthesizing enzymes, which guides processive insertion of septal PG and septum closure^21,22^. In contrast, in the coccus *Staphylococcus aureus* and the ovococcus *Streptococcus pneumoniae*, the septal PG synthetic enzymes also exhibit processive movement around the division ring, but after septal PG synthesis is initiated this movement is driven by PG synthesis itself and not by FtsZ threadmilling^23,25^. A common feature of all these systems is nevertheless a direct link between the dynamics of specific components of the divisome and the rate of PG synthesis and septum closure. Importantly, several cell division inhibitors blocking the constriction process have been shown to directly interact with the divisome in various bacteria^13–15,17^, but the impact of these interactions on the dynamics of the targeted proteins remains unexplored.

Here, we investigated the mechanism of cell division inhibition observed during the development of competence for transformation in the human pathogen *S. pneumoniae* (the pneumococcus). Natural genetic transformation is a conserved mechanism of horizontal gene transfer, which enables the cells to acquire new genetic traits and repair DNA damages in bacteria^26^. It requires that cells enter a differentiation state called competence during which exogenous DNA is imported into the cell and integrated into the chromosome by homologous recombination. In *S. pneumoniae*, competence is transient. It relies on a secreted peptide pheromone, the competence-stimulating peptide (CSP) that spreads through the whole population during exponential phase, and lasts for a short period of time (∼25 min, shorter than the generation time of a single cell)^27–29^. Remarkably, the cell division process is delayed in competent pneumococcal cells, and evidences indicate that this delay contributes to the preservation of genomic integrity in transforming cells^30^. ComM, a membrane protein induced during competence, appears necessary and sufficient to inhibit cell division and to interfere with PG synthesis^30^ This protein also confers immunity to a killing mechanism termed fratricide that can be used by competent cells to acquire DNA from non-competent pneumococci^31–33^. ComM localizes at midcell but does not affect assembly and localization of divisome components, suggesting that it transiently inhibits the active process of constriction. In *S. pneumoniae,* both septal and peripheral (elongation) PG synthesis localize at the site of cell division and occur concomitantly throughout the cell cycle^25,34^. Septal PG synthesis is mediated by the septal FtsW transglycosylase and its PBP2x transpeptidase binding partner, which move together along septal rings^25,34,35^, while the RodA transgycosylase and its cognate PPB2b transpeptidase mediate peripheral PG synthesis and thus elongation of the ovococcal cell^34–37^.

In this study, we sought to investigate the molecular mechanism underlying the ComM-mediated cell division delay of competent cells. We show that ComM is a dynamic protein that co-localizes with and reduces the dynamics of the FtsW:PBP2x complex in *S. pneumoniae*. Furthermore, we provide evidence that reduction of septal PG synthesis may result from ComM-mediated PG precursors shortage.

## Results

### ComM forms discrete, mobile patches around the cell circumference

To understand how ComM interferes with the cell division process during competence, we first examined its dynamics. For this, we constructed a strain containing a functional mNeonGreen-ComM fusion expressed from the native *comM* chromosomal locus under the control of its native competence-induced promoter. The fusion was specifically produced in competent cells with no detectable proteolysis (Extended Data Fig. 1A). Protein levels reached a maximum about 15 minutes after competence induction and then progressively decreased (Extended Data Fig. 1B). The functionality of the fusion was assessed as per its ability to delay cell division and lead to cell elongation during competence (Extended Data Fig. 2A-C)^30,38^. Epifluorescence microscopy confirmed that mNeonGreen-ComM localized in a band at the site of division in competent cells (Extended Data Fig. 3) as previously reported^30^.

To resolve the localization pattern of ComM at the division site with optimal resolution, we vertically immobilized pneumococcal cells in agarose microholes and used highly inclined laminated optical sheet (HILO) microscopy to visualize the entire division plane (Fig 1A). This revealed that ComM forms multiple, independent patches distributed around the cell periphery, which move around the division plane (Fig. 1B, C). Total internal reflection fluorescence microscopy (TIRFM) of cells lying horizontally on the microscope coverslip (Fig. 1D) confirmed directional movements within the ComM ring (Fig. 1E, F). Patches moved in both directions around the cell circumference at an average speed of 12.2 ± 5.3 nm.s^−1^ (Fig. 1G, H). The same speed was measured independently in kymographs around cell circumference of vertically oriented cells (Extended Data Fig. 4). Notably, ComM speed remained constant throughout the development of competence, which lasts for about 20 to 30 minutes in the population^27,29^, and beyond (Fig. 1I), while the number of cells exhibiting a septal signal progressively decreased (Extended Data Fig. 1C).

**Figure 1:**
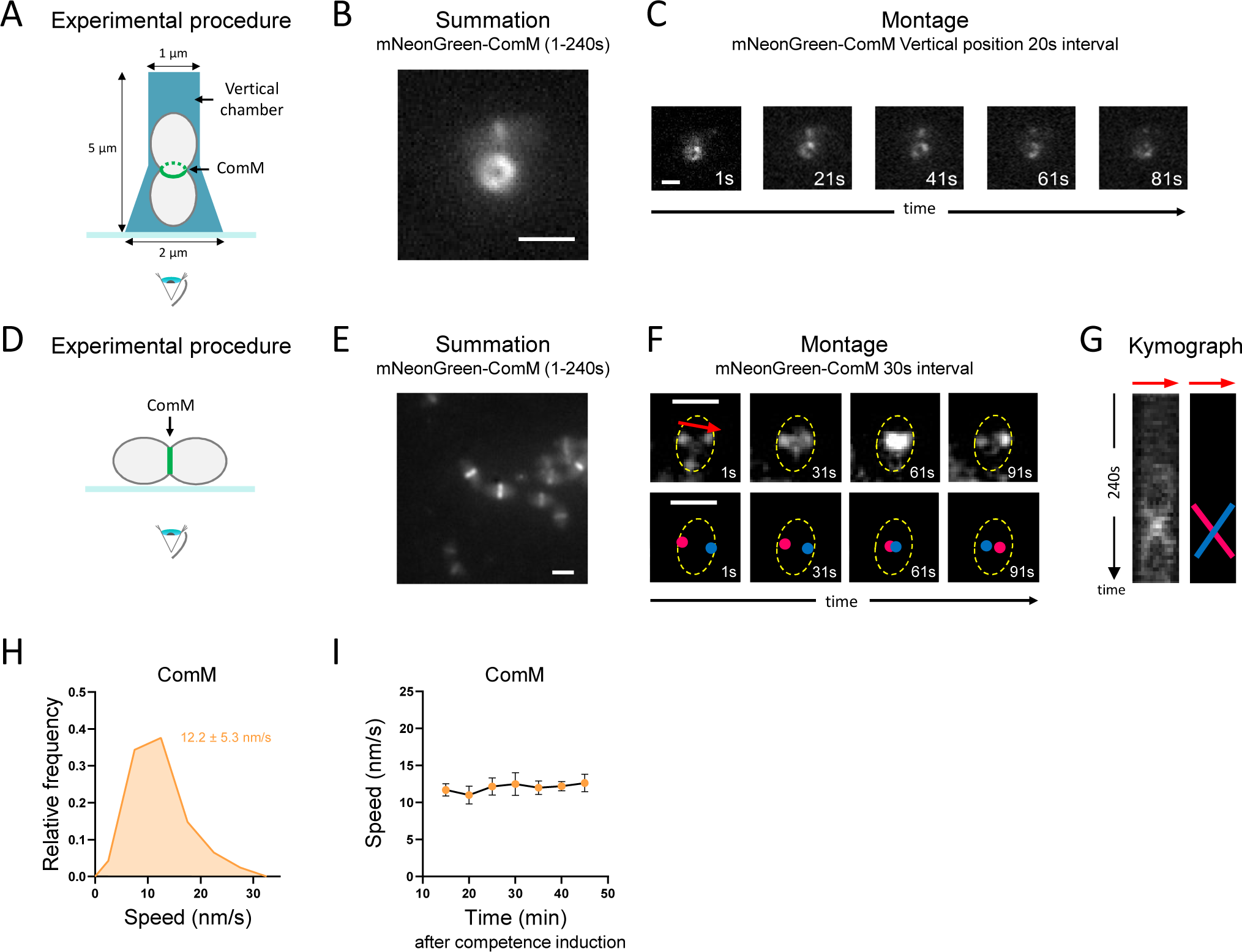
ComM forms dynamic patches moving around the cell circumference. R4601 strain was grown in C+Y medium to early exponential phase and induced to develop competence by CSP addition before imaging. **(A)** Schematics of the method to observe *S. pneumoniae* cells trapped in vertical chambers with the division plane parallel to the coverslip (related to panels *B* and *C*). **(B)** Representative maximum fluorescence intensity projection images. Summation of frames from 240s HILO movies of mNeonGreen-ComM (3s intervals). **(C)** Montage of images at 20s intervals showing mNeonGreen-ComM in vertically oriented cells. **(D)** Schematics of the method to observe *S. pneumoniae* lying horizontally with the division plane orthogonal to the coverslip (related to panels *E* and *F*). **(E)** Representative maximum fluorescence intensity projection images. Summation of frames from 240s TIRFm movies of mNeonGreen-ComM (3s intervals). **(F)** *Upper part*: Montage of TIRFm images at 30s intervals showing mNeonGreen-ComM in cells lying horizontally. The red arrow indicates the trajectory extracted for kymograph analysis (voluntarily slightly shifted to the top to allow the visualization of ComM patches). *Lower part*: Cartoon representation of two ComM patches (red and blue dots). The cell contour is represented in yellow dotted lines. **(G)** Kymograph from 1 to 240s obtained from the ComM trajectory shown in panel *F* (*left*), and its cartoon representation (*right*). The slope of the two ComM patches detected in panel *F* are shown. **(H)** Distribution of speed of mNeonGreen-ComM patches in competent cells (n=752 trajectories). Average speed is indicated. **(I)** mNeonGreen-ComM average speed measured at different time points after competence induction. Error bars represents 95% confidence interval. Representative data are shown from three independent biological replicates. Scale bars, 1 µm.

**Table 1.**
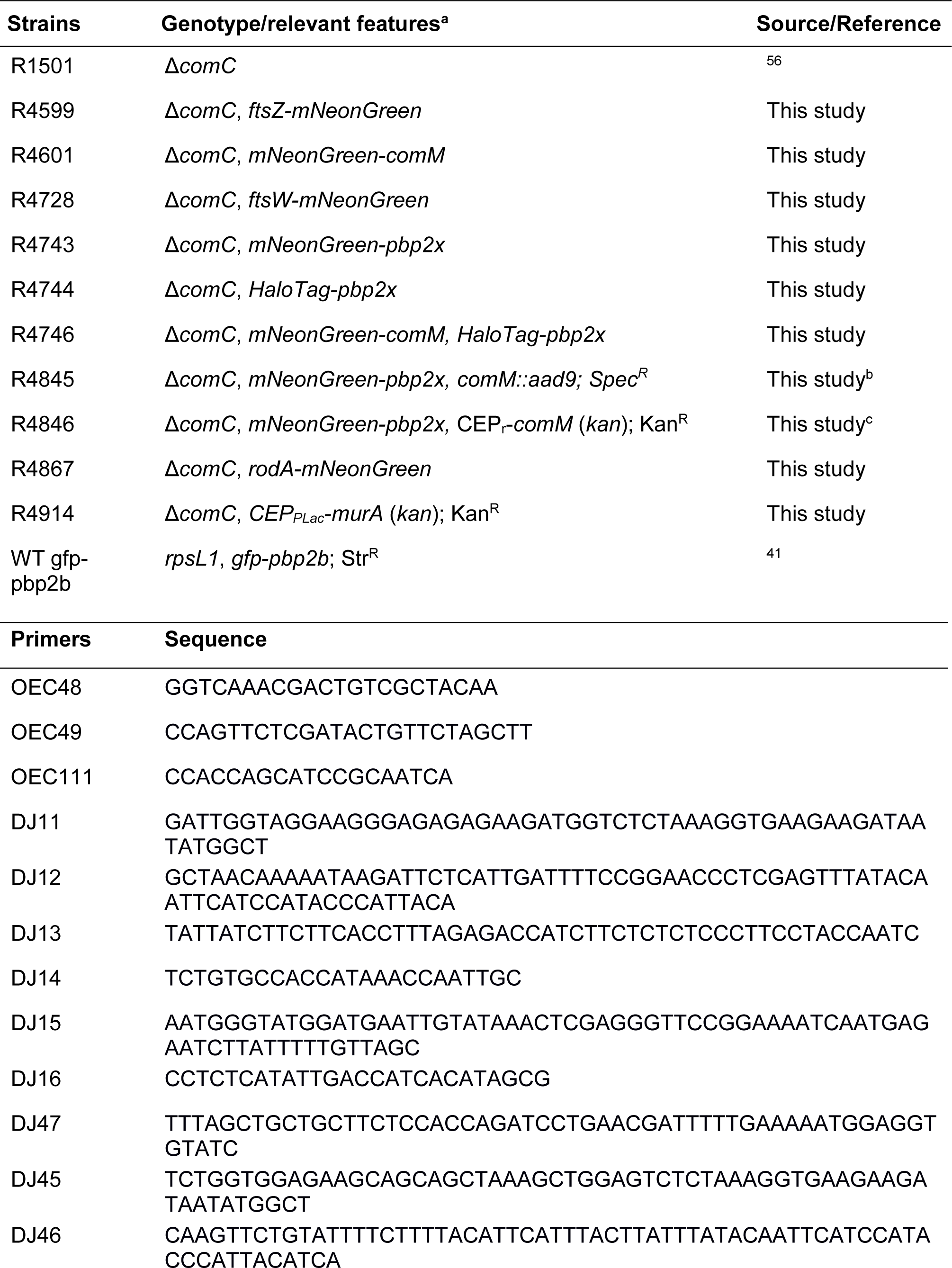
Strains and primers used in this study.

### Dynamics of FtsZ is unaffected during competence

We previously showed that in competent cells, Z-rings assemble at the division site but initiation and completion of the constriction process are transiently inhibited^30^. FtsZ treadmilling dynamics is essential for assembling the divisome^39^. To test whether competence interferes with FtsZ activity, we compared the dynamics of FtsZ in competent and non-competent cells. Cells harboring a functional FtsZ-mNeonGreen fusion expressed from the native locus (Extended Data Fig. 1A, 2D-F and 3) were grown to early exponential phase and induced (or not) to develop competence by adding synthetic CSP to the culture. Demographs showing fluorescence intensity as a function of cell length showed that the deployment of the Z-rings during the cell cycle, from the septa of early divisional cells to the equators of the daughter cells, is unaffected during competence (Extended Data Fig. 5). Maximum projections of TIRFM time-lapses yielded typical patterns of pre-divisional and dividing pneumococcal cells containing mature, equatorial and nascent Z-rings (Fig. 2A), as previously described by the Winkler laboratory^25^. FtsZ rings of early divisional cells become mature septal rings at the onset of division, when most divisome proteins are recruited; then nascent rings form on either side of the mature septal ring and start to move outward to develop into equatorial rings in the future daughter cells at the offset of division (Fig. 2B)^25^. Analysis of the time-lapses confirmed bidirectional movements within the Z-rings (Extended Data Fig. 6A-C), with an average FtsZ filament velocity of about 30 nm.s^−1^ (Fig. 2C), in line with earlier reports in *S. pneumoniae*^25^ and other bacteria^21,22^. Importantly, filament velocity also remained unaffected throughout competence (Fig. 2D). We wondered whether the structure of the Z-rings or the rate of FtsZ treadmilling could be nevertheless impacted by the development of competence at specific stages of the cell cycle (Fig. 2B), which could be masked in our ensemble analysis of the Z-ring population. Precise measurements of the density (total ring intensity) and the speed of FtsZ filaments within five distinct categories of rings with different degrees of maturity (at three different stages of the cell cycle) produced similar values both in non-competent and competent cells at either early or late times of competence (Fig. 2E, F and Extended Data Fig. 6D). Altogether, these results demonstrate that the localization and treadmilling of FtsZ remain unaffected during competence in *S. pneumoniae*.

**Figure 2:**
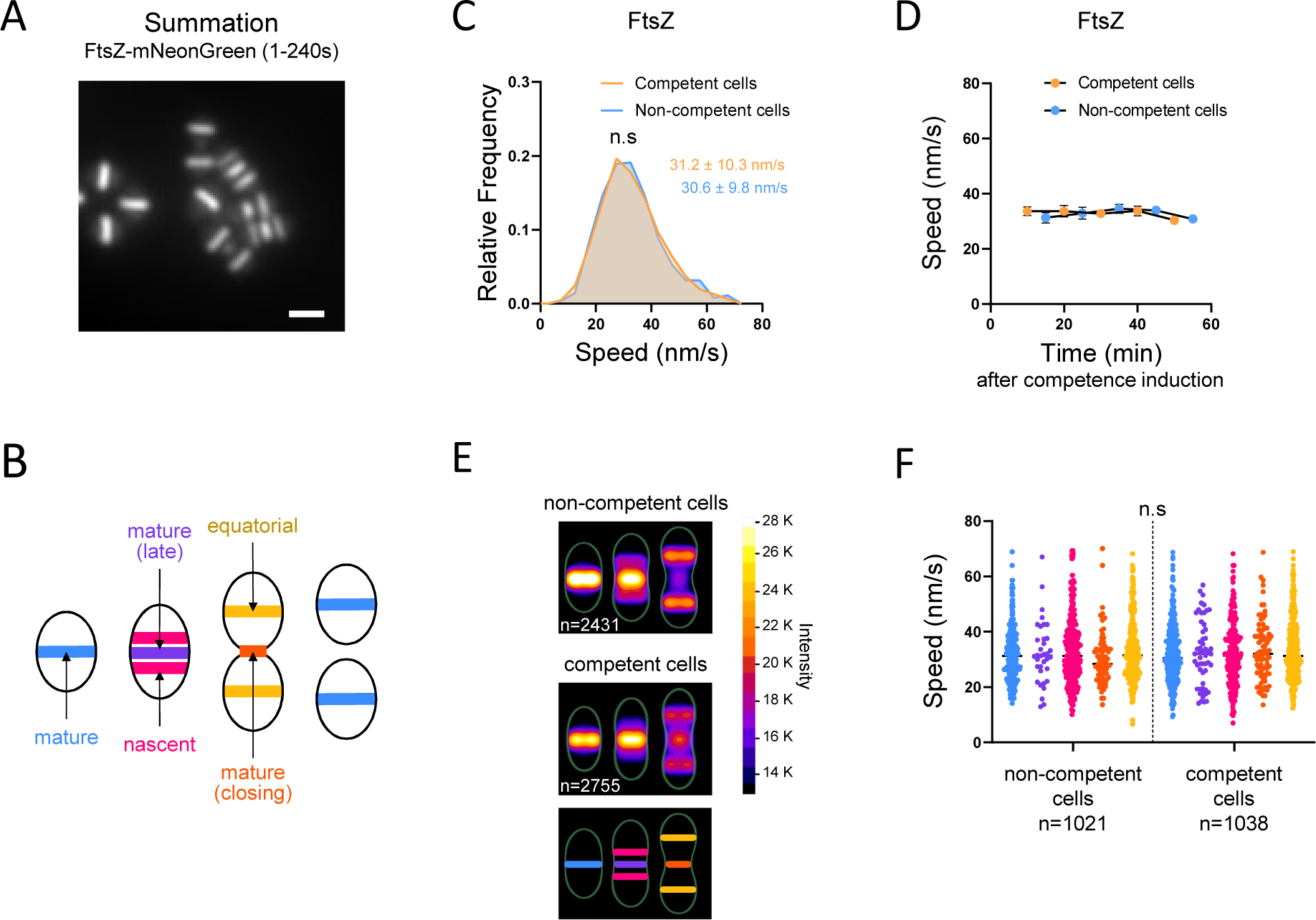
Dynamic of FtsZ is unaffected during competence. Cells (strain R4599) were grown in C+Y medium to early exponential phase and competence was induced, or not, by CSP addition. The data are representative of two independent biological replicates. **(A)** Representative maximum fluorescence intensity projection images. Summation of frames from 240s TIRFm movies of FtsZ-mNeonGreen (3s intervals). Scale bar, 1µm. **(B)** Schematic representation of pneumococcal cells classified into three different classes according to the progression in their cell cycle. Five distinct type of Z-rings are represented. **(C)** Distribution of speed of FtsZ-mNeongreen patches in competent and non-competent cells (922 and 878 trajectories for competent and non-competent cells, respectively). Average speed for each condition is indicated (no significant difference according to the Mann-Whitney nonparametric test). **(D)** FtsZ-mNeonGreen average speed measured at different time points after CSP addition (competent cells) or not (non-competent cells). Error bars represents 95% confidence interval. **(E)** Heatmaps representing the average FtsZ-mNeongreen fluorescence intensity and localization pattern in cells arranged according to size, in non-competent (*up*) and competent (*middle*) cells, and cartoon representation of the corresponding Z-ring types for each cell categories (*down*). N values represent the number of cells analyzed in a single representative experiment. **(F)** Speed recorded in the different Z-ring types in competent and non-competent cells. A total of 1021 and 1038 trajectories (n) were analyzed in non-competent cells and competent cells, respectively. One-way ANOVA statistical analysis showed no significant differences between conditions.

### The mobility of the PBP2x:FtsW complex is reduced in competent cells

As previous labeling experiments of PG synthesis indicated that the rate of PG incorporation at the division site is reduced during competence^30^, we next tested the dynamics of components of the PG synthetic machineries in competent cells. In *S. pneumoniae*, both septal and peripheral PG syntheses, catalyzed by the FtsW:PBP2x and the RodA:PBP2b SEDS-family transglycosylase:bPBP transpeptidase pairs, respectively, occur at the site of cell division and progress separetely^34,35^. We hypothesized that the competence-dependent cell division delay might result from a defect of the septal PG machinery. FtsW and its partner PBP2x have been shown to move along septal rings at the same velocity, which is slower than FtsZ treadmilling and depends of PG synthesis^25^. We constructed functional fusions of FtsW and PBP2x to the mNeonGreen fluorescent protein expressed from the native chromosomal loci. The fusions did not alter growth or cell morphology and showed the typical localization at mature and equatorial rings (Extended Data Fig. 1A, 2D-F and 3). Demographs indicated that the two proteins relocate to the equators of future daughter cells late in the cell cycle, after FtsZ deployment, in both non-competent^25^ and competent cells indistinctly (Extended Data Fig. 5). TIRFM imaging and kymograph analyses showed that in non-competent cells, the mNeonGreen-PBP2x and FtsW-mNeonGreen fusions exhibited bidirectional processive movements at the site of cell division, with average velocities of 19.8 ± 6.2 nm.s^−1^ and 19.8 ± 6.9 nm.s^−1^, respectively (Fig. 3A,B and Extended Data Fig. 7), similar to values reported earlier for halo-tagged PBP2x and FtsW fusions^25^. In addition, FtsW and PBP2x patches exhibited partially diffusive behavior along the membrane (higher background signal in TIRF summations and kymographs relative to FtsZ) (Extended Data Fig. 7), which did not result from cleavage/degradation of the fluorescent fusions (Extended Data Fig. 1A), as previously reported too^25^. Importantly, the average speed of the two fusions was reduced in competent cells (Fig. 3B). We monitored their dynamics as a function of time after CSP addition and found that the velocity of both mNeonGreen-PBP2x and FtsW-mNeonGreen decreased down to about 16 nm.s^−1^ at 10 minutes after competence induction and then gradually recovered, to reach non-competent-like speed 40 to 50 minutes after competence induction (Fig. 3C). We concluded that the development of competence causes a transient deceleration of PBP2x and FtsW motion.

**Figure 3:**
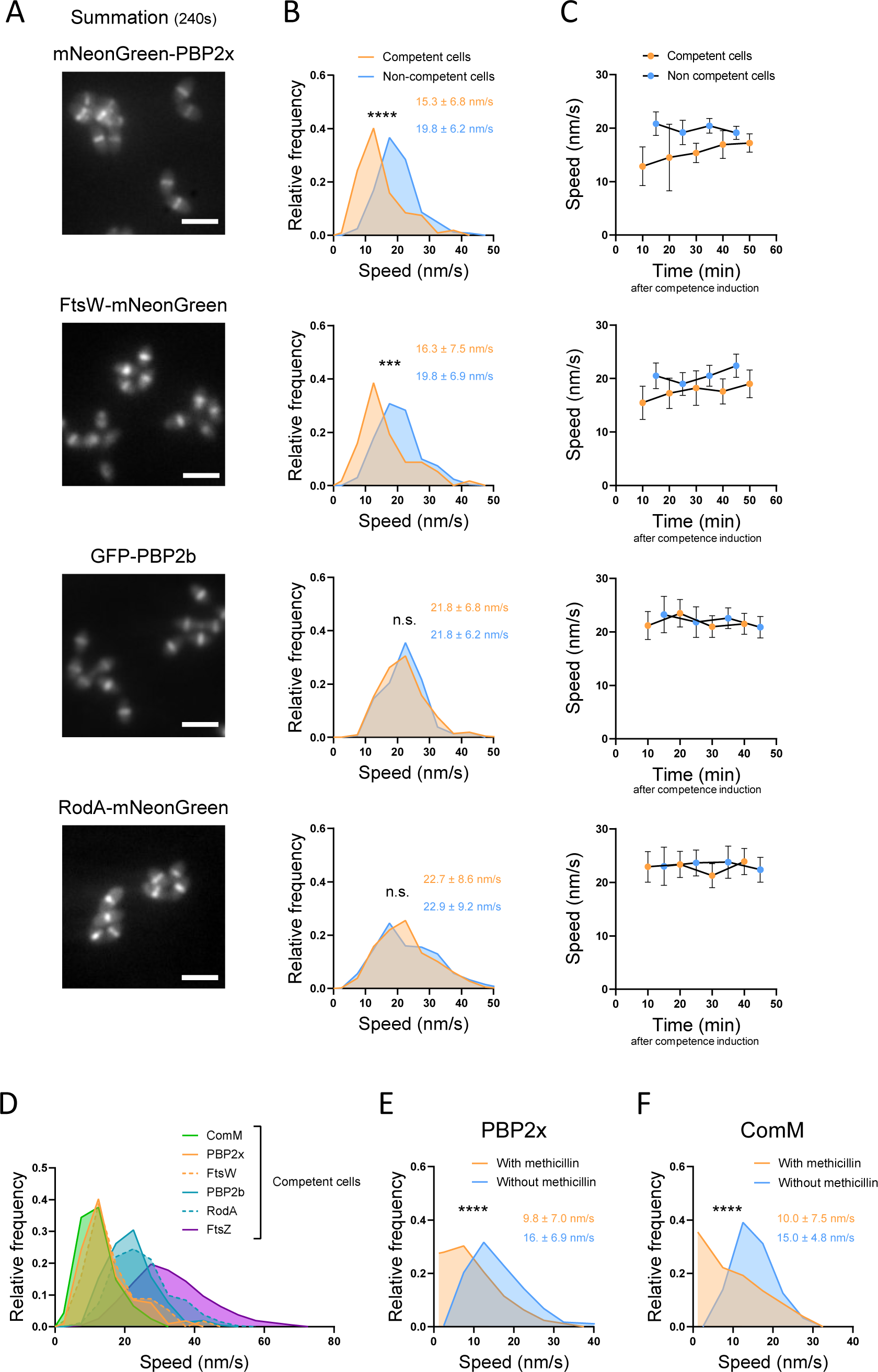
The mobility of PBP2x and FtsW is reduced in competent cells and correlates with ComM. Cells were grown in C+Y medium to early exponential phase and induced to develop competence by CSP addition before imaging by time-lapse TIRF microscopy. Strains analyzed: R4743 (mNeonGreen-PBP2x), R4728 (FtsW-mNeonGreen), WT gfp-pbp2b (GFP-PBP2b), R4867 (RodA-mNeonGreen) and R4599 (FtsZ-mNeonGreen). **(A)** Representative maximum fluorescence intensity projection images. Summation of frames from 240s HILO movies at 3s intervals. Scale bars, 2 µm. **(B)** Distribution of speed in competent and non-competent cells. Average speed is indicated. **(C)** Average speed measured at different time points after competence induction. Error bars represents 95% confidence interval. **(D)** Distribution of speed measured at different time points after competence induction of FtsZ-mNeonGreen, mNeonGreen-PBP2x, FtsW-mNeonGreen, GFP-PBP2b, RodA-mNeonGreen and mNeonGreen-ComM. **(E)** Distribution of speed of mNeongreen-PBP2x patches in competent cells treated, or not, with methicillin. Average speed for each condition is indicated. **(F)** Distribution of speed of mNeonGreen-ComM patches in competent cells treated, or not, with methicillin. Average speed for each condition is indicated. Representative data are shown from two independent biological replicates. A minimum of 150 trajectories were analyzed for each strain and condition. Pairwise comparisons were done with a nonparametric Mann-Whitney test. P values are displayed as follows: ****, P < 0.0001; ***, 0.0001 < P < 0.001; **, 0.001< P < 0.01; *, 0.01 < P < 0.05; ns, P > 0.05.

We then wondered if the peripheral PG synthesis machinery was also affected during competence. RodA and PBP2b have been shown to localize to the site of division in growing *S. pneumoniae* cells^35,40^, but their dynamics has not been investigated so far. Functional fusions of RodA to mNeonGreen (RodA-mNeonGreen) and of PBP2b to GFP (GFP-PBP2b)^41^ expressed from the native locus (Extended Data Fig. 1A, 2D-F and 3) displayed the same deployment pattern from mature to equatorial rings than FtsW and PBP2x in both competent and non-competent cells (Extended Data Fig. 5). TIRFM revealed that RodA and PBP2b also move bidirectionally and processively around the septal rings, albeit at speeds slightly higher than the FtsW:PBP2x complex, with values of about 22 nm.s^−1^ in non-competent cells (Fig. 3 and Extended Data Fig. 7). Importantly, their speed remained unchanged after CSP addition and throughout competence (Fig. 3), suggesting that competence induction has no effect on the peripheral PG machinery. Unperturbed activity of the peripheral PG elongation machinery during competence, when septal PG synthesis is reduced, may explain the increased cell length of competent cells relative to non-competent cells^30^.

Taken together, these findings show that both the bulk of the FtsW:PBP2x and RodA:PBP2b pairs populations exhibit circumferential motions significantly slower than the speed of treadmilling FtsZ filaments along septal rings in either competent (Fig. 3D) and non-competent cells (compare Fig. 2C and 3B). However, their speeds are different, suggesting that the septal and peripheral PG machineries do not move together. This is consistent with the finding that PBP2x and PBP2b localize into separate concentric rings as septation progresses^42^. Furthermore, the speed of the septal FtsW:PBP2x complex but not of the peripheral RodA:PBP2b complex is slowed down during competence, suggesting that septal PG synthesis is reduced and may explain the delay of division in pneumococcal competent cells.

### ComM and PBP2x co-exist in motile nodes at the septal rings of competent cells

We immediately noticed that in competent cells, the FtsW:PBP2x pair slowed down to reach a mean speed similar to that of ComM (Fig. 3D), suggesting that they might share a common motor. FtsW:PBP2x processive movement has been shown to depend on PG synthesis in *S. pneumoniae*^25^. Septal PG synthesis in turn depends on PBP2x transpeptidase activity^44–46^, and PBP2x is required to stimulate FtsW transglycosylase activity^47^. In agreement with this, in the presence of the β-lactam methicillin at a concentration that mainly inhibits PBP2x transpeptidase activity^46,48^, PBP2x motion is virtually arrested in non-competent cells, while FtsZ motion is not (Extended Data Fig. 8)^25^. To test whether the velocity of ComM is correlated with FtsW:PBP2x velocity, we therefore tested the effect of methicillin in competent cells. The velocities of both PBP2x and ComM were drastically reduced in the presence of 0.3 µg/mL methicillin (Fig. 3E, F), suggesting that ComM moves together with FtsW:PBP2x along septal rings. Consistent with this, the kinetics of deployment of ComM from mature to equatorial rings was also similar to that of PBP2x and FtsW (Extended Data Fig. 5). We next generated a strain simultaneously expressing both mNeongreen-ComM and Halo-tag-PBP2x (Extended Data Fig. 1A and 3). Bi-color demographs of this co-localization strain confirmed that the deployment of the two proteins is synchronized during the cell cycle (Fig. 4A). Together, these results indicate that ComM and FtsW:PBP2x exhibit the same dynamics in competent cells.

**Figure 4:**
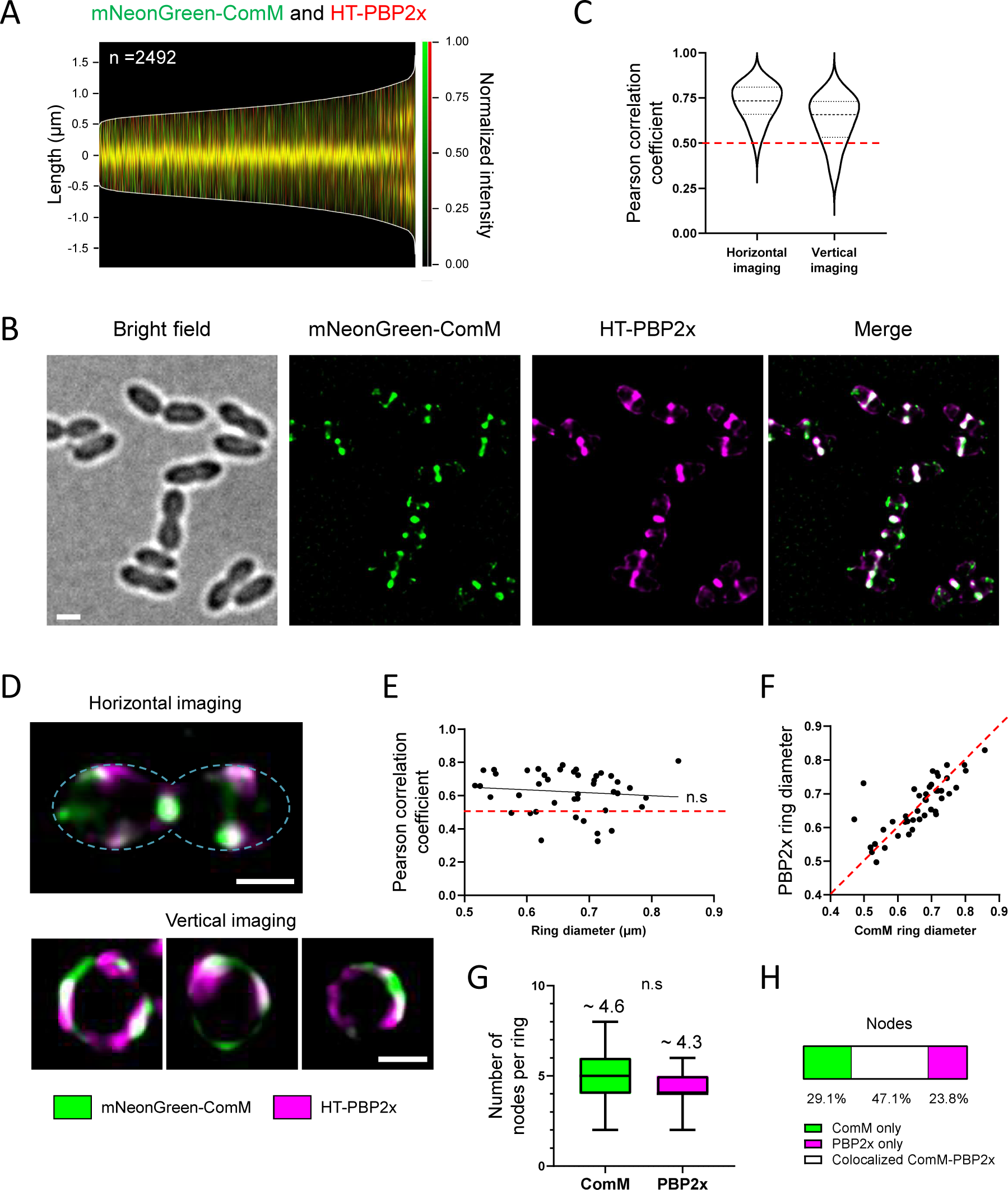
ComM and PBP2x partially colocalize in super resolution microscopy. Strain R4746 containing native mNeonGreen-ComM and HaloTag-PBP2x fusions was grown in C+Y medium to early exponential phase and induced to develop competence by CSP addition for 10 min before imaging. **(A)** Bicolor demograph showing the co-localization signals of mNeonGreen-ComM (green) and HaloTag-PBP2x (red) in competent cells. **(B)** Representative images of bright field fluorescence and merged signals obtained by super resolution SIM² microscopy of cells oriented horizontally. Scale bar is 1 µm. **(C)** Pearson correlation coefficient for a population of cells oriented horizontally (n=100) and vertically (n=43) with PCC=0, no colocalization and PCC=1, total perfect colocalisation. **(D)** Representative images of single cells oriented horizontally (*upper part*) and vertically (*lower part*) obtained by SIM² microscopy. The fluorescence signals of mNeonGreen-ComM and HaloTag-PBP2x are represented in green and magenta, respectively. The white color represents co-localized green and purple signals. Scale bar is 0.5µm. **(E)** PCC in function of the ring diameter (average between ComM and PBP2x ring diameter). Linear regression indicates no correlation between the two parameters. **(F)** PBP2x ring diameter in function of ComM ring diameter. The red dotted reference line corresponds to X=Y (same diameter for each protein). There is no significant difference between the two proteins according to the Mann-Whitney nonparametric test. **(G)** Number of node per ring based on vertical acquisitions SIM² images. Average value for each protein is indicated. There is no significant difference between the two values according to the Mann-Whitney nonparametric test. **(H)** Proportion of the nodes which are fully colocalized (superimposition of a ComM node with a PBP2x node, in white) or not (ComM or PBP2x nodes alone, in green and magenta respectively) in the total of nodes analyzed. The analysis was performed with the support of 100 nm bicolor artificial beads to allow a correct alignment of the channels.

The lateral resolution of TIRFM (∼250 nm) was not sufficient to assess the co-localization of independent patches of ComM and PBP2x at the division rings (Extended Data Fig. 9A). We then moved to super resolution microscopy using Lattice dual iterative SIM (SIM²), which allows imaging at high speed and doubles the conventional resolution of structural illumination microscopy (SIM)^49^. Using SIM² we achieved localizations of mNeonGreen-ComM and Halo-tag-PBP2x with high spatial resolution of ∼70 nm and ∼80 nm, respectively (Extended Data Fig. 9B). No flow through was detected between channels using strains that express the individual fusions (Extended Data Fig. 9C), confirming that these were suitable for co-localization experiments. 2D-SIM^2^ of horizontally immobilized cells demonstrated extensive colocalization of the ComM and PBP2x fusions at the septal rings (Fig. 4B). Pearson’s Correlation Coefficient (PCC) for the fluorescence signals in the two channels confirmed a high level of colocalization (Fig. 4C). Imaging the entire division septum in vertically immobilized cells by 2D-SIM^2^ showed that both proteins form patches (nodes) sparsely distributed around the cell circumference (Fig. 4D and Extended Data Fig. 9D), as previously shown using 3D-SIM^35^, and gave a PCC value similar to that measured in horizontally immobilized cells (Fig. 4C). A similar level of colocalization (PCC) was observed for septal rings of different diameters (Fig. 4E), and ComM and PBP2x rings displayed the same diameter at different division stages (Fig. 4F), suggesting that the two proteins migrate together to the centers of constricting septa. The PBP2x ring was previously shown to separate to the centers of septa (constricting inner ring) at the late stages of cell division whereas the adjacent PBP2b ring (peripheral PG machinery) remains peripheral to it (outer ring)^34,35,45^. Finally, we found that ComM and PBP2x formed a similar number of nodes per septal ring (Fig. 4G) that colocalized in 43% of the cases (Fig. 4H). Taken together, our data provide strong evidence that ComM and PBP2x coexist in motile nodes at the septal ring of competent cells.

### ComM is necessary and sufficient to slow down PBP2x speed

The rate of septal PG synthesis is thought to determine FtsW:PBP2x speed at the septal ring^35^, and ComM, which is required to inhibit cell division, plays an important role in the reduction of the rate of PG synthesis in competent cells^30^. We then wondered if the reduction of speed of FtsW:PBP2x observed during competence depends on ComM. When competence was induced, the *ΔcomM* mutant did not show a delay in cell division (Extended Data Fig. 1A) as expected^30^, and, PBP2x speed did not slow down as in the wild type genetic background (Fig. 5A). We concluded that the reduction of speed of PBP2x observed in competent cells depends on ComM. To test if ComM is sufficient to slowdown PBP2x, we measured PBP2x speed in cells producing ComM outside competence. For this, we used a strain expressing *comM* from an ectopic locus under the control of an inducible promoter^30^. In growing non-competent cells, PBP2x displayed the same average speed than in wild-type cells in the absence of inducer (Fig. 5B, compare to Fig. 3B). However, in the presence of inducer, PBP2x speed was significantly reduced (Fig. 5B). Furthermore, in the presence of inducer the average doubling time and cell length of the population increased, mirroring these two ComM-dependent phenotypes of competent cells (Extended Data Fig. 2A, B)^30^. Taken together, these results indicate that ComM alone is sufficient to slowdown the septal PG synthetic complex during competence.

**Figure 5:**
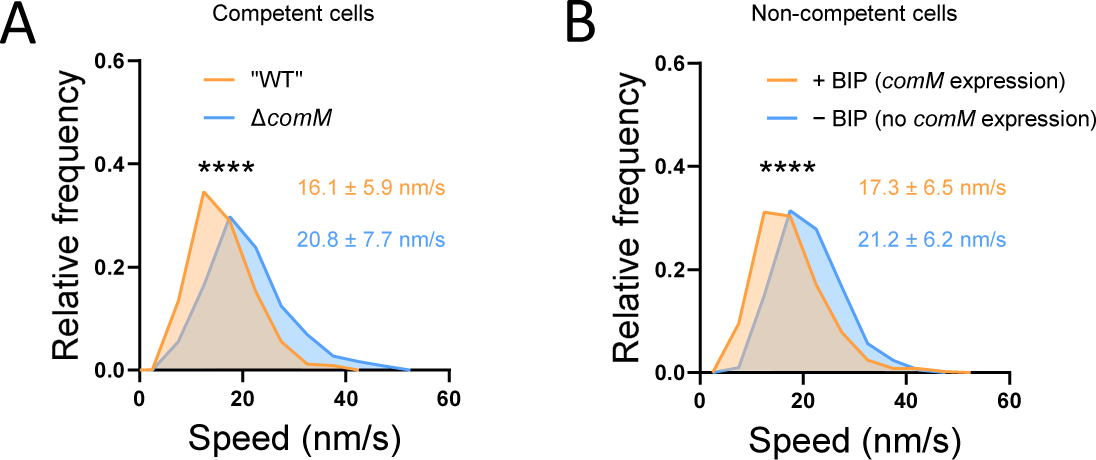
ComM is necessary and sufficient to reduce PBP2x mobility. Cells were grown in C+Y medium to early exponential phase and *comM* expression was induced by addition of CSP (competent cells) or BIP (non-competent cells) before imaging by time-lapse TIRF microscopy. Representative data are shown from two independent biological replicates. **(A)** Distribution of speed of mNeonGreen-PBP2x patches during competence in the Δ*comM* (strain R4845, 538 trajectories) and the wild-type (strain R4743, 644 trajectories) genetic backgrounds. Average speed for each strain is indicated. **(B)** Distribution of speed of mNeonGreen-PBP2x patches in non-competent R4846 cells induced (+ BIP, *comM* expression, 527 trajectories), or not (-BIP, no *comM* expression, 644 trajectories), to produce ComM by BIP induction. Average speed for each condition is indicated. Pairwise comparisons were done with a nonparametric Mann-Whitney test. (****, P < 0.0001).

### ComM may interact with MurA activity to decrease septal PG synthesis

We next sought to investigate the mechanism by which ComM reduces FtsW:PBP2x velocity to delay cell division. Since FtsW:PBP2x velocity is thought to depend on septal PG synthesis rate^25^, ComM must decrease it by reducing the PG synthetic activity of the complex. We found no interaction between ComM and FtsW or PBP2x using the yeast two hybrid system (Fig. 6A), suggesting that ComM does not affect their activity through a direct protein-protein interaction. Interestingly, the velocities of PBP2x and FtsW were found to be reduced by the same amount in a Δ*murZ* mutant background^25^, presumably upon depletion of PG precursors. In *S. pneumoniae*, the first committed step of PG precursor synthesis is catalyzed by two MurA homologues, MurA (SPR1781) and MurZ (SPR0989)^50–52^. They are synthetically lethal but MurZ displays higher catalytic efficiency in vitro^51^, and *murZ* mutant cells, but not *murA* mutants, are larger and display reduced growth^50^. Furthermore, it was recently reported that when the availability of lipid-bound PG precursors is reduced in *S. pneumoniae*, septal PG synthesis is halted while peripheral PG synthesis continues, leading to cell elongation^53^. It was proposed that the activity of the septal PG synthesis enzymes is specifically affected upon PG precursors shortage and outcompeted by the peripheral PG synthesis enzymes^53^. We then wondered if the division delay of competent cells could somehow result from transient depletion of PG precursors via ComM. To explore this possibility, we tested a possible interaction of ComM with several proteins of the PG synthetic pathway using the yeast two hybrid system. Importantly, a specific interaction between ComM and MurA was detected (Fig. 6A). This raised the interesting hypothesis that ComM might inhibit MurA via a direct protein-protein interaction to locally reduce the pool of PG precursors at the division site, reducing the activity of FtsW:PBP2x. In this scenario, artificial overproduction of MurA in competent cells should bypass ComM inhibition and suppress the division delay. When the cellular amounts of MurA were increased by expressing an additional copy of *murA* from the IPTG-inducible promoter P_Lac_, no delay in cell division was detected in competent cells relative to non-competent cells, similarly to what is observed in the *ΔcomM* mutant background (Fig. 6B)^30^. In agreement with this, when MurA was overproduced in competent cells, the velocity of PBP2x was no longer reduced (Fig. 6C). These findings suggest that the specific ComM-MurA interaction detected in the yeast is physiologically relevant, and point towards a model in which ComM reduces FtsW:PBP2x activity via MurA. Further research will be necessary to elucidate the precise molecular mechanism by which ComM reduces septal PG synthesis in competent cells.

**Figure 6:**
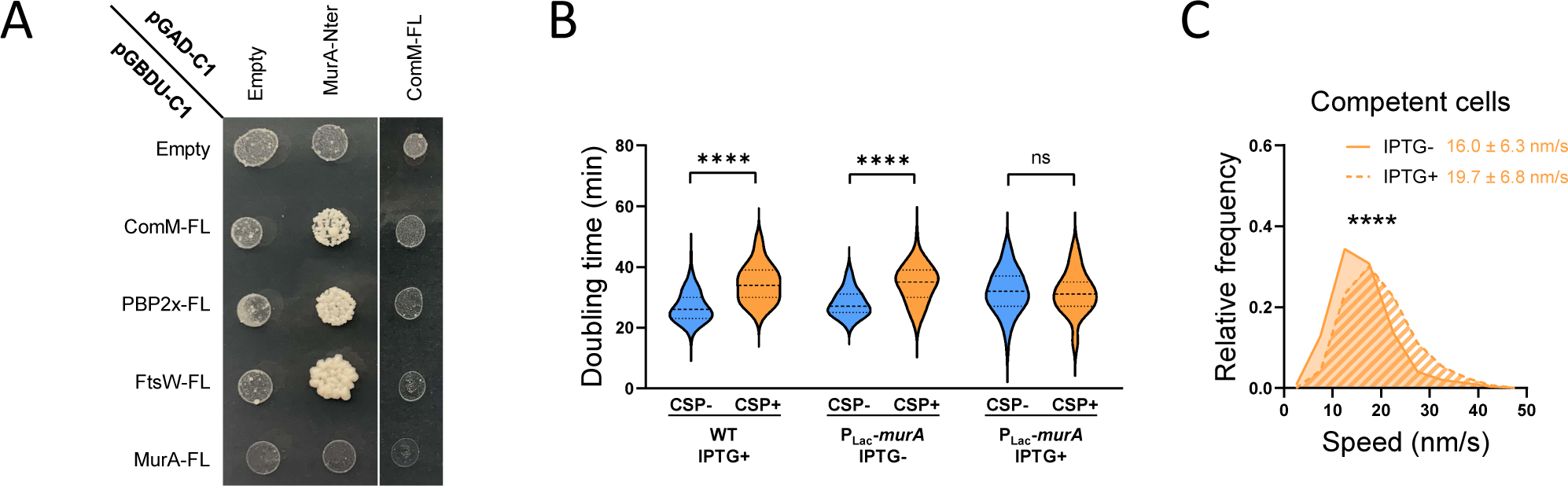
Surproduction of MurA suppress the competence-dependent cell division delay. Cells were grown to early exponential phase in C+Y medium supplemented, or not, with IPTG to induce *murA* expression, and incubated with CSP 10 min before imaging. Representative data are shown from two independent biological replicates. **(A)** Yeast-two-hybrid matrix performed to test interactions between ComM, PBP2x, FtsW and MurA. **(B)** Doubling time of individual cells from strains R1501 (WT) and R4914 (P_lac_-*murA*) based on phase contrast time-lapses analyses. 25th and 75th percentile are shown, with the horizontal line at the median. **(C)** Distribution of speed of mNeonGreen-PBP2x patches in competent cells overexpressing (IPTG+) or not (IPTG-) MurA (strain R4914). Average speed for each condition is indicated. For each condition, a minimum of 250 trajectories were analyzed. Pairwise comparisons were done with a nonparametric Mann-Whitney test. (****, P < 0.0001).

## Discussion

Our results indicate that the transient inhibition of cell division that occurs during competence in *S. pneumoniae* results from ComM-dependent deceleration of FtsW:PBP2x, the SEDS:bPBP pair providing PG polymerase and crosslinking activity in the septal PG synthetic machinery. PG synthesis activity has been shown to power FtsW:PBP2x motion^25^. Thus, a transient deceleration of FtsW:PBP2x in competent cells can explain their reduced septal PG synthesis^30^. We show that ComM forms nodes at the septal rings of competent cells that colocalize partially with and exhibit the same dynamics than PBP2x and FtsW throughout competence. We also show that the SEDS-family polymerase of the PG elongation machinery, RodA, and its cognate bPBP PBP2b exhibit bidirectional processive movement too at the site of cell division, albeit with higher speed than the FtsW:PBP2x pair. The spatial separation of RodA:PBP2b and FtsW:PBP2x in two different concentric rings during septum closure^42^ is therefore accompanied by different speed dynamics, indicating that the two systems function autonomously. Furthermore, while FtsW:PBP2x decelerate during competence, the speed of RodA and PBP2b (thought to reflect the activity of the peripheral PG synthesis machinery), remains unchanged. FtsZ treadmilling also remains unaffected during competence and is faster than the speeds of the two PG synthetic machineries, consistent with what was observed in vegetatively growing *S. pneumoniae* cells^25^. In competent cells, the motion of both ComM and PBP2x was inhibited upon inhibition of PBP2x transpeptidase activity by methicillin, in agreement with the motion of FtsW:PBP2x being driven by PG synthesis^25^ and suggesting that ComM shares the same motor. No direct protein-protein interaction was found between full-length ComM and PBP2x or FtsW in exploratory yeast two hybrid experiments but this does not exclude that a direct interaction may occur in *S. pneumoniae*, or be mediated by a third partner. Further work will be required to investigate possible interactions between ComM and components of the septal PG synthetic machinery allowing ComM motion to be driven by PG synthesis.

Importantly, we show that FtsW:PBP2x deceleration directly depends on ComM and can be suppressed in competent cells together with their division delay by overexpression of MurA, one of the two MurA homologues catalyzing the first committed step of PG precursor synthesis in the cytoplasm. At the overexpression level tested, MurA suppressed the division delay and the reduction of PBP2x speed, but did not rescue the ComM-dependent cell elongation phenotype of competent cells (Extended Data Fig. 10). The septal and elongation machineries must compete for a limited supply of PG precursors in the cell. Although the activity of the septal PG synthesis enzymes may be more specifically affected upon PG precursors shortage^53^, increasing the pool of PG precursors is expected to increase the activity of both the septal and the peripheral PG synthetic machineries, explaining these phenotypes. Thus, it will be interesting to test too the effect of MurA expression on the motion of RodA:PBP2b. Furthermore, the speed of FtsW:PBP2x was found to be reduced in a Δ*murZ* mutant background^25,50^ (note that the effect of *murA* was not tested in this study). MurA is synthetically lethal with MurZ, but MurZ displays higher catalytic efficiency in vitro^51^, and *murZ* mutant cells, but not *murA* mutants, are larger and display reduced growth^50^, suggesting that MurZ has a more prominent role in *S. pneumoniae*. Testing the effect of MurZ overexpression as well as of general PG precursors excess in both the elongation and division machineries, either in competent and non-competent cells, remain questions for future work.

Finally, we found MurA to directly interact with ComM in the yeast. On the basis of the existing literature suggesting that the septal PG machinery is more sensitive to PG precursors shortage^25,53^, through this direct protein-protein interaction ComM might decrease the activity or the levels of MurA, leading to PG precursors shortage that is specifically sensed by the septal PG machinery. Interestingly, the amino acid sequence of ComM possesses one of the three conserved motifs found in the catalytic site of the so-called CAAX proteases (also called the Abi family^54^), which remain uncharacterized in bacteria. Furthermore, mutations in this motif and in several conserved residues that could be part of a CAAX-like catalytic site abolished the immunity of competent cells toward CbpD^33^. This raises the intriguing possibility that ComM might have a protease activity and specifically degrade MurA as substrate. Alternatively, ComM may have lost its proteolytic activity but not the ability to bind its targets and inhibit MurA activity by capturing the protein. However, MurA also displayed a positive interaction with both FtsW and PBP2x in our yeast-two-hybrid experiments. In an alternative scenario, the interaction of ComM with MurA might capture MurA at the division ring and allow its interaction with FtsW:PBP2x to somewhat reduce their activity. Finally, it cannot be excluded that the inhibitory effect of ComM on the activity of the septal PG machinery is independent of MurA, and that overexpression of MurA simply bypasses it by increasing the pool of available PG precursors. ComM could regulate septal PG synthesis through other components of the divisome that directly regulate FtsW:PBP2x activity and/or are involved in the balance between the septal and elongation machineries. Further work will be required to test these hypotheses.

Altogether, our findings provide a better mechanistic understanding of the transient inhibition of cell division that halts cell cycle progression in competent pneumococcal cells and raise intriguing questions about the relative regulatory mechanisms of the different types of PG synthetic machineries. Mechanistic insight provided in this study and future work on the modulation of the activities of the elongasome and the divisome will pave the road toward the discovery of novel ways to disrupt these essential cellular processes for antibiotic development.

## Material and Methods

### General methods

*S. pneumoniae* strains and oligonucleotide primers used in this study are listed in Supplementary Tables 1–3. *S. pneumoniae* strains were all constructed in the R1501 background, which is derived from strain R800^55^. This strain contains the *ΔcomC* mutation and cannot develop competence spontaneously^56^. Stock cultures were routinely grown at 37°C to OD550∼0.3 in Todd-Hewitt medium (BD Diagnostic System) supplemented with 0.5% Yeast Extract (THY) or C+Y medium^57^; after addition of 15% (vol/vol) glycerol, stocks were kept frozen at −70°C. For the monitoring of growth, pre-cultures grown in C+Y medium to OD_550_∼0.3 were diluted 1 in 40 in C+Y medium and further grown to OD_550_ ∼ 0.1. Cells were then diluted 40-folds in C+Y medium and distributed into a 96-well microplate (300 µL per well). OD_492_ values were recorded throughout incubation at 37°C in a Varioskan Flash luminometer (Thermo Fisher ScientificWaltham, MA, USA). Note that we measured OD at 492nm in the Varioskan luminoter, which gives similar results to monitor cell growth as measurements at OD 550nm. Transformation was performed as described previously^58^. Antibiotic concentrations (μg ml^−1^) used for the selection of *S. pneumonia*e transformants were: kanamycin (Kan), 250; spectinomycin (Spec), 100; and streptomycin (Str), 200. Ectopic expression of *comM* (strain R)and *murA* (strain) were achieved by adding 250 μg ml^−1^ BIP-1^59,60^ and 100µM IPTG, respectively, to the cultures

### Strain constructions

Strain R4599, containing a *ftsZ*-*mNeonGreen* fusion at the *ftsZ* endogeneous locus, was generated as follows. Primers were designed to amplify PCR products containing: (I) the upstream region and the entire coding sequence of the *ftsZ* (*spr1510*) gene (2044 bp; oligonucleotides OEC48 and DJ47, and R1501 DNA as template); (II) a 12 amino-acid linker, L5^61^, and the *orf* of the *mNeonGreen* gene (770 bp; oligonucleotides DJ45 and DJ46, and plasmid DNA pPEPY-Plac-Link-mNeonGreen^62^ as template); and (III) the downstream region of the *ftsZ* gene (2048 bp; oligonucleotides DJ48 and OEC49, and R1501 DNA as template). The PCR products were gel-purified and used as templates in a SOEing PCR using the outer primers OEC48 and OEC49. The resulting SOEing PCR product was subsequently used to transform strain R1501 without selection, as previously described^63^.

Strain R4601, harboring a *mNeonGreen* fusion at the *comM* endogeneous locus, was created as follows. Primers were designed to amplify PCR products containing: (I) the upstream region of the *comM* (*spr1762*) gene (2049 bp; oligonucleotides DJ14 and DJ13, and R1501 DNA as template); (II) the *orf* of the *mNeonGreen* gene followed by a 6 amino-acid linker, L1^41^, yielding a 776 bp PCR fragment (oligonucleotides DJ11 and DJ12, and plasmid DNA pPEPY-Plac-Link-mNeonGreen^62^ as template); and (III) the *comM* orf and its downstream region (2154 bp; oligonucleotides DJ15 and DJ16, and R1501 DNA as template). The PCR products were gel-purified and used as templates in a SOEing PCR using the outer primers DJ14 and DJ16. The resulting SOEing PCR product was subsequently used to transform strain R1501 without selection.

Strain R4743, carrying a *mNeonGreen-pbp2x* fusion, was created as follows. Primers were designed to amplify PCR products containing: (I) the upstream region of the *pbp2x* (*spr0304*) gene (2111 bp; oligonucleotides OCN450 and OCN451, and R1501 DNA as template); (II) the *orf* of the *mNeonGreen* gene and the L1 linker^41^ yielding a 776 bp PCR fragment (oligonucleotides OCN452 and OCN453, and plasmid DNA pPEPY-Plac-Link-mNeonGreen^62^ as template); and (III) the *pbp2x* orf and its downstream region (2128 bp; oligonucleotides OCN454 and OCN455, and R1501 DNA as template). The PCR products were gel-purified and used as templates in a SOEing PCR using the outer primers OCN450 and OCN455. The resulting SOEing PCR product was subsequently used to transform strain R1501 without selection.

Strain R4744, containing a *HaloTag-pbp2x* fusion, was created as follows. Primers were designed to amplify PCR products containing: (I) the upstream region of the *pbp2x* gene (2100 bp; oligonucleotides OCN450 and OCN456, and R1501 DNA as template); (II) a 963 bp PCR fragment carrying the *orf* encoding the HaloTag sequence and a 15 amino-acids linker, L6^25^ (oligonucleotides OCN457 and OCN458, and a DNA fragment containing the HaloTag sequence generated by Integrated DNA Technologies, see below); and (III) the *pbp2x* orf and its downstream region (2127 bp; oligonucleotides OCN459 and OCN455, and R1501 DNA as template). The PCR products were gel-purified and used as templates in a SOEing PCR using the outer primers OCN450 and OCN455. The resulting SOEing PCR product was subsequently used to transform strain R1501 without selection.

Strain R4728, containing a *ftsW*-*mNeonGreen* fusion at the *ftsW* endogeneous locus, was generated as follows. Primers were designed to amplify PCR products containing: (I) the upstream region and the entire coding sequence of the *ftsW* (*spr0973*) gene (2043 bp; oligonucleotides OCN440 and OCN441, and R1501 DNA as template); (II) the L5 linker^61^ and the *orf* of the *mNeonGreen* gene (804 bp; oligonucleotides OCN442 and OCN443, and plasmid DNA pPEPY-Plac-Link-mNeonGreen^62^ as template); and (III) the downstream region of the *ftsW* gene (2168 bp; oligonucleotides OCN444 and OCN445, and R1501 DNA as template). The PCR products were gel-purified and used as templates in a SOEing PCR using the outer primers OEC440 and OEC445. The resulting SOEing PCR product was subsequently used to transform strain R1501 without selection, as previously described^63^.

Strain R4867 harboring a *rodA*-*mNeonGreen* fusion was generated as follows. Primers were designed to amplify PCR products containing: (I) the upstream region and the coding sequence of the *rodA* (*spr0712*) gene (2163 bp; oligonucleotides DJ59 and DJ60, and R1501 DNA as template); (II) the L5 linker^61^ and the *orf* of the *mNeonGreen* gene (798 bp; oligonucleotides DJ61 and DJ62, and plasmid DNA pPEPY-Plac-Link-mNeonGreen^62^ as template); and (III) the downstream region of the *rodA* gene (2128 bp; oligonucleotides DJ63 and DJ64, and R1501 DNA as template). The PCR products were gel-purified and used as templates in a SOEing PCR using the outer primers DJ59 and DJ64. The resulting SOEing PCR product was subsequently used to transform strain R1501 without selection.

To over-produce MurA, we constructed a strain, R4914 (P_Lac_-*murA*), allowing ectopic expression of *murA* (*murA1*, *spr1781*^50,52^) at the CEP chromosomal expression platform^64^ under the control of the IPTG-inducible P_Lac_ promoter^65^. Primers were designed to amplify PCR products containing: (I) the 5’ region of the CEP platform including the *lacI* gene and the P_Lac_ promoter (3783 bp; oligonucleotides OCN564 and OCN568, and R3310^65^ DNA as template); (II) the *orf* of the *murA* gene (1324 bp; oligonucleotides OCN569 and OCN570, and R1501 DNA as template); and (III) the 3’ region of the CEP platform including the gene conferring resistance to kanamycine (2899 bp; oligonucleotides OCN571 and OEC111, and R3310 DNA as template). The PCR products were gel-purified and used as templates in a SOEing PCR using the outer primers OCN564 and OEC111. The resulting SOEing PCR product was subsequently used to transform strain R1501 and clones resisting to kanamycin were selected. All constructs generated by PCR were confirmed by DNA sequencing.

The DNA fragment containing the HaloTag sequence was purchased from Integrated DNA Technologies (IDT). This fragment carries the HaloTag sequence codon optimized for *S. pneumoniae* published with the Winkler laboratory^25^ with a few modifications. It also contains the L5 (bold) and L6 (underlined) linkers at the 5’ and 3’ extremities, respectively: **TCAGGATCTGGTGGAGAAGCAGCAGCTAAAGCTGGA**GCTGAAATTGGTACTGGTTTTCCATTTGATCCACATTATGTTGAAGTTTTGGGTGAACGTATGCATTATGTTGATGTTGGTCCACGTGATGGTACTCCAGTTTTGTTTTTGCATGGTAATCCAACAAGTTCTTATGTTTGGCGAAATATTATTCCACATGTTGCTCCAACACATCGTTGTATTGCTCCAGATCTTATTGGTATGGGTAAATCTGATAAACCAGATTTGGGTTATTTTTTTGATGATCATGTTCGTTTTATGGATGCTTTTATTGAAGCTTTGGGTTTGGAAGAAGTTGTTTTGGTTATTCATGATTGGGGTTCTGCTTTGGGTTTTCATTGGGCTAAACGTAATCCAGAACGTGTTAAAGGTATTGCTTTTATGGAATTTATTCGTCCAATTCCAACATGGGATGAATGGCCAGAATTTGCTCGTGAAACATTTCAAGCTTTTCGTACAACAGATGTTGGTCGTAAATTGATTATTGATCAAAATGTTTTTATTGAAGGTACATTGCCAATGGGTGTTGTTCGTCCATTGACAGAAGTTGAAATGGATCATTATCGTGAACCATTTTTGAATCCAGTTGATCGTGAACCATTGTGGCGTTTTCCAAATGAATTGCCAATCGCTGGTGAACCAGCTAATATTGTTGCTTTGGTTGAAGAATATATGGATTGGTTGCATCAATCTCCAGTTCCAAAATTGTTGTTTTGGGGTACACCAGGTGTTTTGATTCCACCAGCTGAAGCTGCTCGTTTGGCTAAATCTTTGCCAAATTGTAAAGCTGTTGATATTGGTCCAGGTTTGAATTTGTTGCAAGAAGATAATCCAGATTTGATTGGTTCTGAAATTGCTCGTTGGTTGTCTACATTGGAAATCTCTGGT<u>TTGGAAGGATCAGGACAAGGACCAGGAAGTGGTCAAGGTTCAGGT</u>

### Whole-cell extracts preparation and immunoblot analysis

Pneumococcal cells were grown in C+Y medium at 37°C to exponential phase (OD550 ∼0.1) and treated or not with CSP for 15 min, and samples (3 ml) were collected by centrifugation. Cell pellets were stored at −80°C. Whole-cell extracts were prepared by resuspension of cell pellets in 50 µl lysis buffer [10 mM Tris pH 8.0, 1 mM EDTA, 0.01% (wt/vol) DOC, 0.02% (wt/vol) SDS] and incubation at 37°C for 10 min followed by addition of 50 µl loading buffer [0.25 M Tris pH 6.8, 6% (wt/vol) SDS, 10 mM EDTA, 20% (vol/vol) Glycerol] containing 10% (vol/vol) β-mercaptoethanol. Samples were heated for 15 min at 50°C prior to loading.

For immunoblot analysis, whole cell extracts samples were loaded on Biorad mini-PROTEAN TGX stain-free 4–15% pre-cast gels. Gels were activated by ultraviolet (UV) exposure for 45 s using a Bio-Rad ChemiDoc MP imager to visualize and estimate total protein per lane. Proteins were then transferred to a nitrocellulose membrane, using a Turbo Blot transfer unit (Bio-Rad). mNeonGreen and Halotag fusion proteins were detected using polyclonal anti-mNeonGreen (NC, unpublished) and anti-Halotag (Promega) antibodies diluted at 1:10,000 and 1:2,000 respectively. Primary antibodies were detected using peroxidase-conjugated goat anti-rabbit immunoglobulin G (Sigma) diluted 1:10,000. Membranes were further incubated with clarity chemiluminescence substrate (Bio-Rad), and imaged on the ChemiDoc MP. Detection was performed using Image Lab 5.2 software (Bio-Rad).

### Sample preparation for microscopy

After gentle thawing of stock cultures, aliquots were inoculated at OD_550_ 0.006 in C+Y medium and grown at 37°C to an OD_550_ of 0.3. These precultures were inoculated (1/50) in C+Y medium and incubated at 37°C to an OD_550_ of 0.1. Then, competence was induced with synthetic CSP1 (50 ng ml^−1^). Ectopic *comM* expression was induced by 250 µg ml^−1^ BIP-1^59,60^. 0.8 μl of this suspension was immobilized on a 1.2% C+Y agarose-coated microscope slide. For time-lapse microscopy experiments, cells grown at 37°C to an OD_550_ of 0.1, were induced with CSP1 (50 ng ml^−1^) for 5 minutes and 0.8 µl was immediately spotted on a microscope slide containing a slab of 1.2% C+Y agarose and covered with a cover glass (# 1.5) before imaging.

### Vertical cell mounting

For vertical imaging, silicon molds containing pyramid-shaped pillars (1µm in diameter at the top, 2µm at the base and 5µm high) were prepared for inverse replication (technological support of LAAS-CNRS micro and nanotechnology platform, Toulouse, France). Agarose pads were prepared by pouring melted 6% agarose in C+Y medium on slides, subsequently covered with the silica molds to generate microholes. 6 µL of fresh cells were deposited on agarose pads and the slides were centrifuged in a mini-centrifuge (10 s, 2000 g). 10 µl of 1% agarose in C+Y medium were then poured on top of the agarose pad to immobilize cells in the microholes, and covered with a cover glass before imaging.

### Epifluorecence, TIRF and HILO microscopy

All TIRF and HILO time-lapses were acquired on an automated Zeiss Elyra PS1 microscope with climate control chamber equipped with a 100×/1.46 NA Apochromat oil immersion objective. Laser excitation at 488 nm and 561 nm was set between 10 and 15% of the maximum excitation power and with exposure time of 100-150ms depending of the strain. Images were collected with an emCCD camera (Andor iXon), using electron-multiplying gains (300×) settings attached to a 2.5x magnification lens. Acquisition was controlled by the Zen Software (Zeiss) software package. Epifluorescence snapshots were performed on a previously described setup^66^ equipped with a 100 x objective and with laser excitation at 488 nm and 561 nm set at 30% of the maximum excitation power with exposure time of 500-750 ms depending of the strain.

### Velocity measurements of dynamic proteins

Time-lapses image sequences were stabilized using the Fiji plugin “Image stabilizer” in translation mode with a maximum pyramid levels of 1, a template update coefficient of 0.90, 200 maximal iterations and an error tolerance of 0.0000001. Velocities of directionally moving structures were quantified in image time series using a script implemented in MATLAB (Mathworks, R2018b). This method automatically generates kymographs distributed at regular intervals (∼1 pixel) perpendicular to a manually defined axis (segment connecting the two opposite poles of the bacteria). The kymographs obtained are aligned side-by-side to generate a single 2D image (the vertical axis corresponds to time and the horizontal axis represents spatial displacement). Since directionally moving structures appear as a tilted line in a kymograph, velocity is quantified by measuring the slope of the line.

### Velocity measurement of mNeonGreen-ComM using circular kymographs

The circular kymographs were generated in Fiji^67^ using the ‘Multi Kymograph’ plugin. First, a circle was manually drawn on FtsZ rings (centered and of the same diameter). The selection was then transformed into a polygonal line to generate kymographs with a 3-pixel linewidth. Finally, the velocities were quantified by measuring the slopes on the kymographs.

### Demograph analysis

Demographs showing protein fluorescence intensity as a function of cell length were generated by using Microbe J (version 5.13n)^68^ with the following parameters: (area [μm2] 0.6-20; length [μm] 0.5-max; width ([μm] 0.5-2; circularity [0-1] 0-max; curvature [0-max] 0-max; sinuosity [0-max] 0-max; angularity [rad] 0-0.25; solidity [0-max] 0.85-max; intensity [0-max] 0-max). Cells with regular shapes and sizes but excluded from analyses due to close proximity to other cells were manually added back to analyses. In addition, late-divisional cells were verified to correctly count them as two separated cells if necessary as previously described^25^. The demographs were normalized as proposed in microbeJ.

### Heat map analysis

Heat map showing protein fluorescence average intensity in the cell were generated by using Microbe J (version 5.13n)^68^ with the following parameters as criteria to define the categoires of cells: A: [SHAPE.length/SHAPE.width]<=1.7; B: [SHAPE.length/SHAPE.width]>1.7 and [SHAPE.length]<=1.6; C: [SHAPE.length/SHAPE.width]>1.7 and [SHAPE.length]>1.6 D: [SHAPE.length/SHAPE.width]>2 and [SHAPE.circularity]<0.8; E: [SHAPE.length/SHAPE.width]> 2 and [SHAPE.circularity]<0.7.

### Lattice SIM imaging and SIM² image reconstruction

Lattice SIM imaging was conducted mainly as previously described^69^ using an Elyra 7 AxioObserver (Zeiss) inverted microscope. Here, we used a 63×/NA 1.46 objective (Zeiss, alpha Plan-Apochromat 63x/1.46 Oil Korr M27) yielding a final pixel size of 64.5 nm for raw images. Fluorescence was excited using a 488 nm (100 mW) and a 561 nm (100 mW) laser lines at 80% and 40% of maximal output power for 488 and 561 laser respectively. Exposure time per phase was 100ms and 200ms for 488 nm and 561 nm laser respectively. Temperature was maintained at 30°C during acquisitions.

Lattice SIM image reconstruction was performed via Zen Software (Zeiss, black edition) using the nonlinear iterative reconstruction algorithm SIM² with 2D+ processing active. General settings were set to “Live” and “Strong” with the following specifics: 20 iterations, a regularization weight of 0.02, a processing sampling of 4×, and an output sampling of 4×. Advanced filter parameters were set to “Best fit” and “Median” with a sectioning value of 100 and Baseline “Yes”. Following SIM² reconstruction, all images had a lateral pixel size of 16.1 nm. The Fiji plugin “TurboReg” was used to performed XY-alignment between the separate fluorescent channels. The plugin was set to “Rigid body”, in “fast” quality and automatic mode and used 100 nm bicolor artificial beads located within the same field of view. PCC was calculated on individual rings with the Fiji plugin “JaCoP” v2.1.121.

### Single cell doubling time and morphologic quantification

Phase contrast time-lapses image sequences were stabilized using the Fiji plugin “Image stabilizer” as described above for velocity measurements. Segmentation of phase-contrast images was performed using the image segmentation tool Omnipose^70^. Cell morphology (cell width and length) was estimated as the width and length of the minimum area rectangle around the segmented mask. Doubling times were measured from tracks of individual cells generated with the overlap tracker in Trackmate^71^. Tracking graphs were extracted and analyzed using a custom Python script that worked as follows: a division was detected as a node (representing a cell) having 2 children instead of one. A low number of artefacts, caused mostly by misdetection of dead elements in the background as cells, were removed from analysis by removing branches of the graph with less than 5 nodes. Division time for a cell was defined as the difference between the time it divided and the time it was created by division. Because of the short (1h) acquisition time and to avoid biasing results towards low division times we considered only divisions occurring in cells created in the first 20 minutes.

### Yeast two hybrid

Gene-coding sequences of *S. pneumoniae* selected proteins (ComM, PBP2x, FtsW, MurA (Spr1781) and the N-terminal domain of MurA, MurA-Nter) were PCR amplified using R1501 DNA as template, and inserted in Gal4-based plasmids, PGAD-C1 and PGBD-C1^72^, using the In-Fusion HD cloning kit (Clontech). Primers used were OCN334 and OCN430 (ComM), OCN364 and OCN365 (FtsW) OCN366 and OCN367 (PBP2x), OCN390 and OCN391 (MurA), OCN390 and OCN392 (MurA-Nter), and OCN324 and OCN325 (PGAD-C1 and PGBD-C1).

Resulting plasmids were then established in the yeast strains PJ69-4a and PJ69-4α. *Saccharomyces cerevisiae* cells expressing *S. pneumoniae* proteins as GAL4 BD fusions were mated with cells expressing some of these proteins as GAL4 AD fusions. Binary interactions were revealed by growth of diploid cells after 7 days at 30°C on synthetic complete medium^73^ lacking leucine, uracil, and either adenine (to select for expression of the ADE2 interaction reporter) or histidine (to select for expression of the HIS3 interaction reporter). Controls with empty vector plasmids (*i.e.*, carrying only the BD or AD domain) were systematically included.

### Statistical analyses

All statistical analyses were performed with Prism 10 (GraphPad Software, LLC). Pairwise comparison between two conditions were done with a nonparametric Mann-Whitney test, P values are displayed as follows: ****, 0.0001; ***, 0.0001; **, 0.001; *,0.01; ns, 0.05.

### Data availability

All relevant data and strains supporting the findings of the study are available in this article and its Supplementary information files, or from the corresponding authors on reasonable request.

### Code availability

The analysis code used in this study was written in Python and is deposited on GitHub at https://github.com/aurelien-barbotin/dl_for_mic

## Acknowledgments

This work is part of DJ’s PhD thesis, co-supervised by N.C and R.C.-L. We thank Jan willem veening for his mNeonGreen plasmid and Christophe Grangeasse for the WT GFP-PBP2B strain. We also thank Anne-Lise Soulet (LMGM) and Léa Wagner (Micalis institute) for their experimental assistance. Silicon molds containing pyramid-shaped pillars for vertical imaging were produced by the LAAS-CNRS micro and nanotechnologies platform member of the French RENATECH network. This work was funded by the Centre National de la Recherche Scientifique, Université Paul Sabatier, and grants from the Agence Nationale de la Recherche (EXStasis, ANR-17-CE13-0031 to P.P., N.C. and R.C.-L.) and from the European Research Council (ERC) under the Horizon 2020 research and innovation program (grant agreement No 772178 to R.C.-L.), and by fellowship ‘Fin de thèse’ from the Fondation pour la Recherche Médicale (FDT202204014744 to D.J.).

## Author contributions

N.C. and R.C.-L. conceived the project. D.J., A.L., I.M-B. and N.C. performed experiments. D.J generated fluorescent strains, performed fluorescence microscopy experiments (TIRFM and SIM^2^) and yeast-two-hybrid experiments. A.L. assisted in SIM^2^ experiments. I.M-B. generated the plasmids for yeast-two hybrid experiments. N.C. generated overexpression and fluorescent pneumococcal strains, and performed western blot analysis. D.J. C.B., A.B. and A.L. processed, analyzed and interpreted microscopy data. C.B. established the TIRFM workflow, circular kymographs and assisted in general microscopy data collection and analysis. A.B. wrote the python scripts that measured doubling time and cell morphology in microscopy timelapses. P.P/N.C. and R.C.-L. provided infrastructure and scientific advice. R.C.-L. provided the TIRFM and SIM microscopes. D.J., N.C. and R.C.-L. wrote the manuscript.

## Competing interests

The authors declare no competing interests.

## Notes

### Competing Interest Statement

The authors have declared no competing interest.

## References

1. Heinrich, K., Leslie, D. J. & Jonas, K. Modulation of bacterial proliferation as a survival strategy. Adv Appl Microbiol 92, 127–171 (2015).

2. Jonas, K. To divide or not to divide: control of the bacterial cell cycle by environmental cues. Curr Opin Microbiol 18, 54–60 (2014).

3. Handler, A. A., Lim, J. E. & Losick, R. Peptide inhibitor of cytokinesis during sporulation in Bacillus subtilis. Mol Microbiol 68, 588–599 (2008).

4. Valladares, A., Velázquez-Suárez, C. & Herrero, A. Interactions of PatA with the Divisome during Heterocyst Differentiation in Anabaena. mSphere 5, e00188–20 (2020).

5. Xing, W.-Y., Liu, J., Zhang, J.-Y., Zeng, X. & Zhang, C.-C. A proteolytic pathway coordinates cell division and heterocyst differentiation in the cyanobacterium Anabaena sp. PCC 7120. Proc Natl Acad Sci U S A 119, e2207963119 (2022).

6. Del Sol, R., Pitman, A., Herron, P. & Dyson, P. The product of a developmental gene, crgA, that coordinates reproductive growth in Streptomyces belongs to a novel family of small actinomycete-specific proteins. J Bacteriol 185, 6678–6685 (2003).

7. Del Sol, R., Mullins, J. G. L., Grantcharova, N., Flärdh, K. & Dyson, P. Influence of CrgA on assembly of the cell division protein FtsZ during development of Streptomyces coelicolor. J Bacteriol 188, 1540–1550 (2006).

8. Flärdh, K. & Buttner, M. J. Streptomyces morphogenetics: dissecting differentiation in a filamentous bacterium. Nat Rev Microbiol 7, 36–49 (2009).

9. Huisman, O. & D’Ari, R. An inducible DNA replication-cell division coupling mechanism in E. coli. Nature 290, 797–799 (1981).

10. Mukherjee, A., Cao, C. & Lutkenhaus, J. Inhibition of FtsZ polymerization by SulA, an inhibitor of septation in Escherichia coli. Proc Natl Acad Sci U S A 95, 2885–2890 (1998).

11. den Blaauwen, T., Hamoen, L. W. & Levin, P. A. The divisome at 25: the road ahead. Current Opinion in Microbiology 36, 85–94 (2017).

12. Bisson-Filho, A. W. et al. FtsZ filament capping by MciZ, a developmental regulator of bacterial division. Proc Natl Acad Sci U S A 112, E2130–2138 (2015).

13. Masser, E. A. et al. DNA damage checkpoint activation affects peptidoglycan synthesis and late divisome components in *Bacillus subtilis*. Molecular Microbiology 116, 707–722 (2021).

14. Modell, J. W., Hopkins, A. C. & Laub, M. T. A DNA damage checkpoint in Caulobacter crescentus inhibits cell division through a direct interaction with FtsW. Genes Dev 25, 1328–1343 (2011).

15. Modell, J. W., Kambara, T. K., Perchuk, B. S. & Laub, M. T. A DNA damage-induced, SOS-independent checkpoint regulates cell division in Caulobacter crescentus. PLoS Biol 12, e1001977 (2014).

16. Bojer, M. S., Frees, D. & Ingmer, H. SosA in Staphylococci: an addition to the paradigm of membrane-localized, SOS-induced cell division inhibition in bacteria. Curr Genet 66, 495–499 (2020).

17. Kim, H. K. & Harshey, R. M. A Diguanylate Cyclase Acts as a Cell Division Inhibitor in a Two-Step Response to Reductive and Envelope Stresses. mBio 7, e00822–16 (2016).

18. Burby, P. E. & Simmons, L. A. Regulation of Cell Division in Bacteria by Monitoring Genome Integrity and DNA Replication Status. J Bacteriol 202, e00408–19 (2020).

19. Bojer, M. S. et al. SosA inhibits cell division in *Staphylococcus aureus* in response to DNA damage. Mol Microbiol 112, 1116–1130 (2019).

20. Briggs, N. S., Bruce, K. E., Naskar, S., Winkler, M. E. & Roper, D. I. The Pneumococcal Divisome: Dynamic Control of Streptococcus pneumoniae Cell Division. Front. Microbiol. 12, 737396 (2021).

21. Bisson-Filho, A. W. et al. Treadmilling by FtsZ filaments drives peptidoglycan synthesis and bacterial cell division. Science 355, 739–743 (2017).

22. Yang, X. et al. GTPase activity–coupled treadmilling of the bacterial tubulin FtsZ organizes septal cell wall synthesis. Science 355, 744–747 (2017).

23. Monteiro, J. M. et al. Peptidoglycan synthesis drives an FtsZ-treadmilling-independent step of cytokinesis. Nature 554, 528–532 (2018).

24. Li, Y. et al. MapZ Forms a Stable Ring Structure That Acts As a Nanotrack for FtsZ Treadmilling in *Streptococcus mutans*. ACS Nano 12, 6137–6146 (2018).

25. Perez, A. J. et al. Movement dynamics of divisome proteins and PBP2x:FtsW in cells of *Streptococcus pneumoniae*. Proc Natl Acad Sci USA 116, 3211–3220 (2019).

26. Johnston, C., Martin, B., Fichant, G., Polard, P. & Claverys, J.-P. Bacterial transformation: distribution, shared mechanisms and divergent control. Nat Rev Microbiol 12, 181–196 (2014).

27. Håvarstein, L. S., Coomaraswamy, G. & Morrison, D. A. An unmodified heptadecapeptide pheromone induces competence for genetic transformation in Streptococcus pneumoniae. Proc Natl Acad Sci U S A 92, 11140–11144 (1995).

28. Prudhomme, M., Berge, M., Martin, B. & Polard, P. Pneumococcal Competence Coordination Relies on a Cell-Contact Sensing Mechanism. PLoS Genet 12, e1006113 (2016).

29. Alloing, G., Martin, B., Granadel, C. & Claverys, J. P. Development of competence in Streptococcus pneumonaie: pheromone autoinduction and control of quorum sensing by the oligopeptide permease. Mol Microbiol 29, 75–83 (1998).

30. Bergé, M. J. et al. A programmed cell division delay preserves genome integrity during natural genetic transformation in Streptococcus pneumoniae. Nat Commun 8, 1621 (2017).

31. Guiral, S., Mitchell, T. J., Martin, B. & Claverys, J.-P. From The Cover: Competence-programmed predation of noncompetent cells in the human pathogen Streptococcus pneumoniae: Genetic requirements. Proceedings of the National Academy of Sciences 102, 8710–8715 (2005).

32. Havarstein, L. S., Martin, B., Johnsborg, O., Granadel, C. & Claverys, J.-P. New insights into the pneumococcal fratricide: relationship to clumping and identification of a novel immunity factor. Mol Microbiol 59, 1297–1037 (2006).

33. Straume, D., Stamsås, G. A., Salehian, Z. & Håvarstein, L. S. Overexpression of the fratricide immunity protein ComM leads to growth inhibition and morphological abnormalities in Streptococcus pneumoniae. Microbiology (Reading) 163, 9–21 (2017).

34. Trouve, J. et al. Nanoscale dynamics of peptidoglycan assembly during the cell cycle of Streptococcus pneumoniae. Current Biology 31, 2844–2856.e6 (2021).

35. Perez, A. J. et al. Organization of peptidoglycan synthesis in nodes and separate rings at different stages of cell division of *Streptococcus pneumoniae*. Mol. Microbiol. 115, 1152–1169 (2021).

36. Philippe, J., Vernet, T. & Zapun, A. The elongation of ovococci. Microb Drug Resist 20, 215–221 (2014).

37. Berg, K. H., Stamsås, G. A., Straume, D. & Håvarstein, L. S. Effects of low PBP2b levels on cell morphology and peptidoglycan composition in Streptococcus pneumoniae R6. J Bacteriol 195, 4342–4354 (2013).

38. Mortier-Barrière, I., Polard, P. & Campo, N. Direct Visualization of Horizontal Gene Transfer by Transformation in Live Pneumococcal Cells Using Microfluidics. Genes 11, 675 (2020).

39. Bisson-Filho, A. W. et al. Treadmilling by FtsZ filaments drives peptidoglycan synthesis and bacterial cell division. Science 355, 739–743 (2017).

40. Morlot, C., Zapun, A., Dideberg, O. & Vernet, T. Growth and division of Streptococcus pneumoniae: localization of the high molecular weight penicillin-binding proteins during the cell cycle. Mol Microbiol 50, 845–855 (2003).

41. Fleurie, A. et al. Interplay of the serine/threonine-kinase StkP and the paralogs DivIVA and GpsB in pneumococcal cell elongation and division. PLoS Genet 10, e1004275 (2014).

42. Perez, A. J. et al. Organization of peptidoglycan synthesis in nodes and separate rings at different stages of cell division of *Streptococcus pneumoniae*. Mol. Microbiol. 115, 1152–1169 (2021).

43. Yang, X. et al. A two-track model for the spatiotemporal coordination of bacterial septal cell wall synthesis revealed by single-molecule imaging of FtsW. Nat Microbiol 6, 584–593 (2021).

44. Peters, K. et al. Streptococcus pneumoniae PBP2x mid-cell localization requires the C-terminal PASTA domains and is essential for cell shape maintenance. Mol Microbiol 92, 733–755 (2014).

45. Tsui, H.-C. T. et al. Pbp2x localizes separately from Pbp2b and other peptidoglycan synthesis proteins during later stages of cell division of Streptococcus pneumoniae D39. Mol Microbiol 94, 21–40 (2014).

46. Land, A. D. et al. Requirement of essential Pbp2x and GpsB for septal ring closure in Streptococcus pneumoniae D39. Mol Microbiol 90, 939–955 (2013).

47. Taguchi, A. et al. FtsW is a peptidoglycan polymerase that is functional only in complex with its cognate penicillin-binding protein. Nat Microbiol 4, 587–594 (2019).

48. Kocaoglu, O., Tsui, H.-C. T., Winkler, M. E. & Carlson, E. E. Profiling of β-lactam selectivity for penicillin-binding proteins in Streptococcus pneumoniae D39. Antimicrob Agents Chemother 59, 3548–3555 (2015).

49. Giacomelli, G. et al. Subcellular Dynamics of a Conserved Bacterial Polar Scaffold Protein. Genes (Basel) 13, 278 (2022).

50. Tsui, H.-C. T. et al. Negative regulation of MurZ and MurA underlies the essentiality of GpsB- and StkP-mediated protein phosphorylation in Streptococcus pneumoniae D39. Mol Microbiol 120, 351–383 (2023).

51. Du, W. et al. Two active forms of UDP-N-acetylglucosamine enolpyruvyl transferase in gram-positive bacteria. J Bacteriol 182, 4146–4152 (2000).

52. Hoskins, J. et al. Genome of the bacterium Streptococcus pneumoniae strain R6. J Bacteriol 183, 5709–5717 (2001).

53. Dewachter, L. et al. Amoxicillin-resistant Streptococcus pneumoniae can be resensitized by targeting the mevalonate pathway as indicated by sCRilecs-seq. Elife 11, e75607 (2022).

54. Kjos, M., Snipen, L., Salehian, Z., Nes, I. F. & Diep, D. B. The abi proteins and their involvement in bacteriocin self-immunity. J Bacteriol 192, 2068–2076 (2010).

55. Lefevre, J. C., Claverys, J. P. & Sicard, A. M. Donor deoxyribonucleic acid length and marker effect in pneumococcal transformation. J Bacteriol 138, 80–86 (1979).

56. Dagkessamanskaia, A. et al. Interconnection of competence, stress and CiaR regulons in Streptococcus pneumoniae: competence triggers stationary phase autolysis of ciaR mutant cells: Competence, stress and autolysis in S. pneumoniae. Molecular Microbiology 51, 1071–1086 (2004).

57. Mortier-Barrière, I., Campo, N., Bergé, M. A., Prudhomme, M. & Polard, P. Natural Genetic Transformation: A Direct Route to Easy Insertion of Chimeric Genes into the Pneumococcal Chromosome. in Streptococcus pneumoniae (ed. Iovino, F.) vol. 1968 63–78 (Springer New York, 2019).

58. Martin, B., Prudhomme, M., Alloing, G., Granadel, C. & Claverys, J.-P. Cross-regulation of competence pheromone production and export in the early control of transformation in Streptococcus pneumoniae. Mol Microbiol 38, 867–878 (2000).

59. de Saizieu, A. et al. Microarray-based identification of a novel Streptococcus pneumoniae regulon controlled by an autoinduced peptide. J Bacteriol 182, 4696–4703 (2000).

60. Reichmann, P. & Hakenbeck, R. Allelic variation in a peptide-inducible two-component system of Streptococcus pneumoniae. FEMS Microbiol Lett 190, 231–236 (2000).

61. van Raaphorst, R., Kjos, M. & Veening, J.-W. Chromosome segregation drives division site selection in *Streptococcus pneumoniae*. Proc Natl Acad Sci USA 114, E5959–E5968 (2017).

62. Keller, L. E., Rueff, A.-S., Kurushima, J. & Veening, J.-W. Three New Integration Vectors and Fluorescent Proteins for Use in the Opportunistic Human Pathogen Streptococcus pneumoniae. Genes 10, 394 (2019).

63. Quevillon-Cheruel, S. et al. Structure-function analysis of pneumococcal DprA protein reveals that dimerization is crucial for loading RecA recombinase onto DNA during transformation. Proc Natl Acad Sci U S A 109, E2466–2475 (2012).

64. Guiral, S. et al. Construction and evaluation of a chromosomal expression platform (CEP) for ectopic, maltose-driven gene expression in Streptococcus pneumoniae. Microbiology (Reading) 152, 343–349 (2006).

65. Johnston, C., Mortier-Barriere, I., Khemici, V. & Polard, P. Fine-tuning cellular levels of DprA ensures transformant fitness in the human pathogen Streptococcus pneumoniae. Mol Microbiol 109, 663–675 (2018).

66. Billaudeau, C. et al. Contrasting mechanisms of growth in two model rod-shaped bacteria. Nat Commun 8, 15370 (2017).

67. Schindelin, J., et al. Fiji: an open-source platform for biological-image analysis. Nat Methods 9, 676–682 (2012).

68. Ducret, A., Quardokus, E. M. & Brun, Y. V. MicrobeJ, a tool for high throughput bacterial cell detection and quantitative analysis. Nat Microbiol 1, 16077 (2016).

69. Lablaine, A. et al. A new fluorescence-based approach for direct visualization of coat formation during sporulation in Bacillus cereus. Sci Rep 13, 15136 (2023).

70. Cutler, K. J. et al. Omnipose: a high-precision morphology-independent solution for bacterial cell segmentation. Nat Methods 19, 1438–1448 (2022).

71. Ershov, D. et al. TrackMate 7: integrating state-of-the-art segmentation algorithms into tracking pipelines. Nat Methods 19, 829–832 (2022).

72. James, P., Halladay, J. & Craig, E. A. Genomic libraries and a host strain designed for highly efficient two-hybrid selection in yeast. Genetics 144, 1425–1436 (1996).

73. Guide to yeast genetics and molecular and cell biology. A,1,1998. (Academic Pr, 1998).

