## Supplemental Figures for "Transient inhibition of cell division in competent pneumococcal cells results from deceleration of the septal peptidoglycan complex"

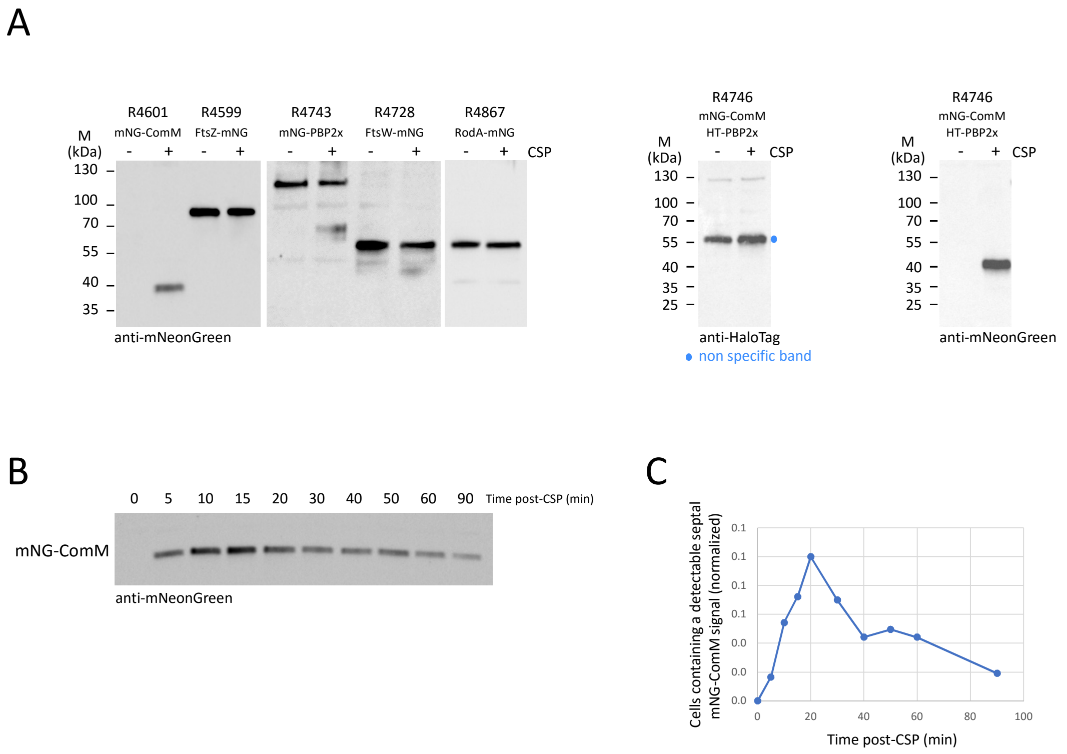


**Extended Data Fig. 1: The strains are fully functional.**

Representative data are shown from two or three independent biological replicates.

**(A)** Western blots with anti mNeonGreen and anti-HaloTag antibodies, on the strains of this study. Cells were induced to develop competence for 15 minutes. Whole cell extracts were prepared and analyzed by immunoblot using anti-mNeonGreen and anti-HaloTag antibodies. Blue dot indicates a non-specific band detected by the anti-HaloTag antibodies.

**(B)** Western blot detection of mNeonGreen-ComM at different times after competence induction (post-CSP). Whole cell extracts were analyzed with anti-mNeonGreen antibodies.

**(C)** Percentage of cells containing a detectable mNeonGreen-ComM signal at the septum, at different times after competence induction (post-CSP).

Extended Data Fig. 2

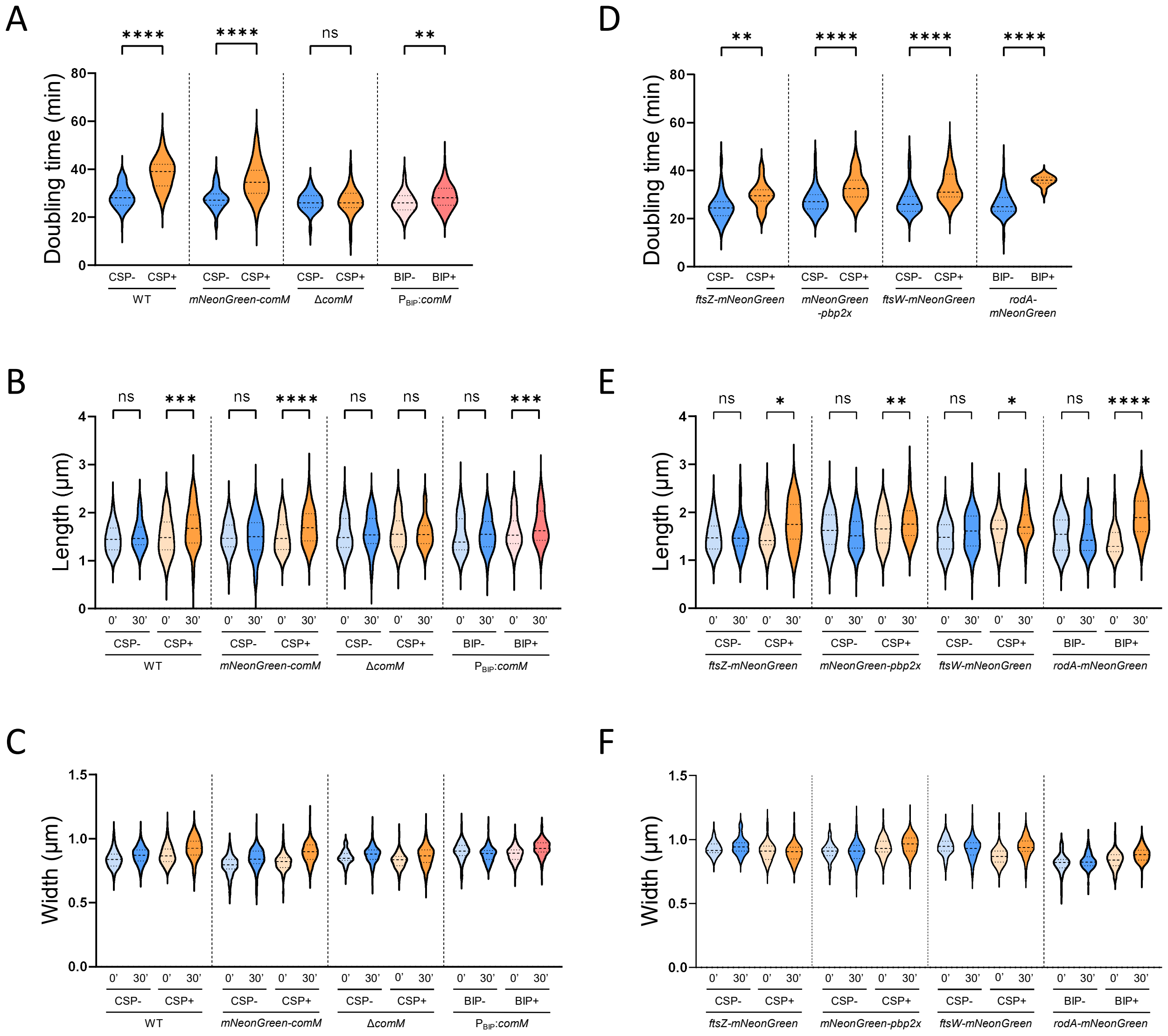


**Extended Data Fig. 2: Functionality of the fluorescent fusions used in this study**
Strains containing native mNeonGreen fusions of ComM (strain R4601), FtsZ (strain R4599), PBP2x (strain R4743), FtsW (strain R4728), RodA (strain R4867), as well as native mNeonGreen fusion of ComM combined with HaloTag fusion of PBP2x (strain R4746), are expressed and have similar doubling time and cell dimensions compared to wild-type cells (strain R1501). Cells were grown in C+Y medium to early exponential phase and competence was induced (+) or not (-) by CSP addition.
**(A,D)** Doubling time of single cells based on phase contrast time-lapses analyses (see Material and Methods). 25th and 75th percentile are shown, with the horizontal line at the median.
**(B-F)** Cell length (B,E) and cell width (C,F) distributions measured on phase contrast microscopy images using > 100 cells for each strain. Cells were incubated with CSP or BIP for 30 minutes before imaging. 25th and 75th percentile are shown, with the horizontal line at the median.
Representative data from three independent biological replicates.

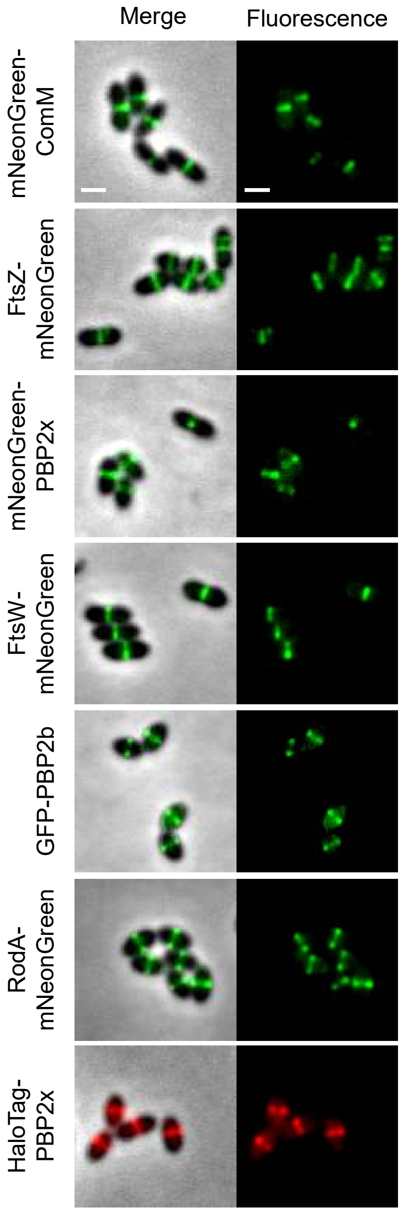


**Extended Data Fig. 3: ComM, FtsZ, PBP2x, FtsW, PBP2b and RodA localize to the division ring at midcell**. Strains used in this study were analyzed by epifluorescence microscopy. Representative fluorescence and merged images between phase-contrast (gray) and mNeonGreen or GFP (green), or Janelia fluor HaloTag ligand (red) signals are shown. Scale bars, 1µm.

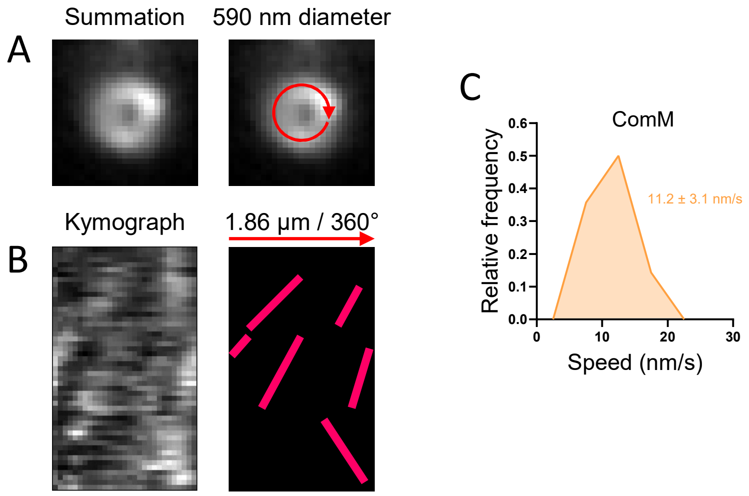


**Extended Data Fig. 4: ComM forms multiple dynamic patches moving in both directions around the cell circumference**.

R4601 cells were grown in C+Y medium to early exponential phase and induced to develop competence by CSP addition for 10 min before imaging. Cells were immobilized vertically and imaged by time-lapse HILO microscopy at 3s intervals. Representative data are shown from two independent biological replicates.

**(A)** Representative maximum fluorescence intensity projection images. Summation of frames from 240s HILO movie of mNeonGreen-ComM.

**(B)** Radial kymograph generated following the red line shown in A (*left*); Cartoon version of the kymograph with multiple trajectories shown in red.

**(C)** Distribution of speed of mNeonGreen-ComM patches in competent cells (n=16 trajectories). Average speed is indicated.

### Extended Data Fig. 5

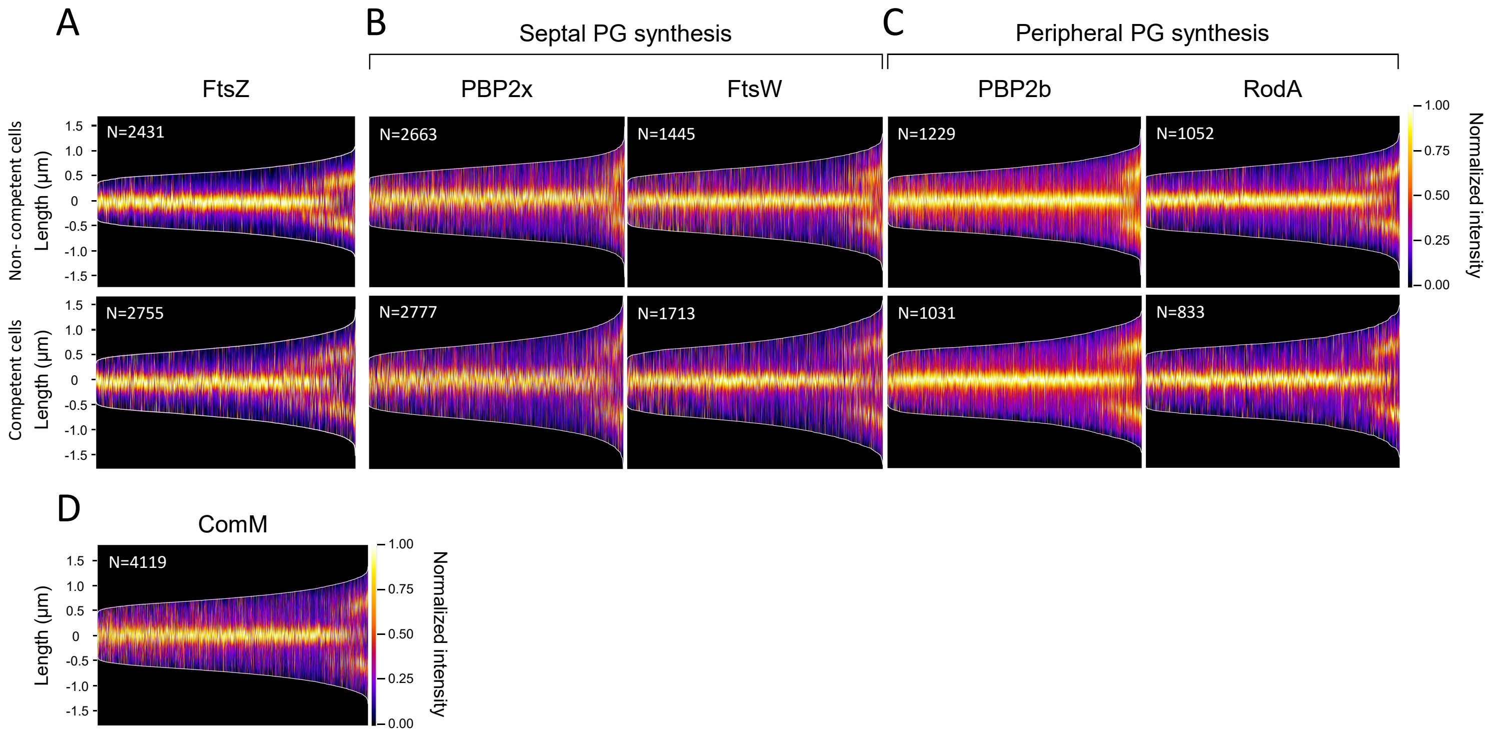


**Extended Data Fig. 5: Deployment of Divisome proteins in competent and non-competent cells.**
Cells were grown in C+Y medium to early exponential phase and competence was induced (+) or not (-) by CSP addition for 30 minutes before imaging. The data are representative of three independent biological replicates. Cells were ordered by cell length and represented by a heatmap. The number of cells analyzed is indicated (N).

(A-E) Demographs showing the localization signal of FtsZ-mNeongreen (R4599) (A), proteins of the septal PG synthesis machinery mNeongreen-PBP2x (R4743), and FtsW-mNeongreen (R4728) (B), proteins of the peripheral PG synthesis machinery GFP-PBP2b and RodA-mNeongreen (R4867) (C) in non-competent (*upper panel*) and competent cells (*lower panel*).

(D) Demograph showing the localization signal of mNeongreen-ComM (R4601) in competent cells.

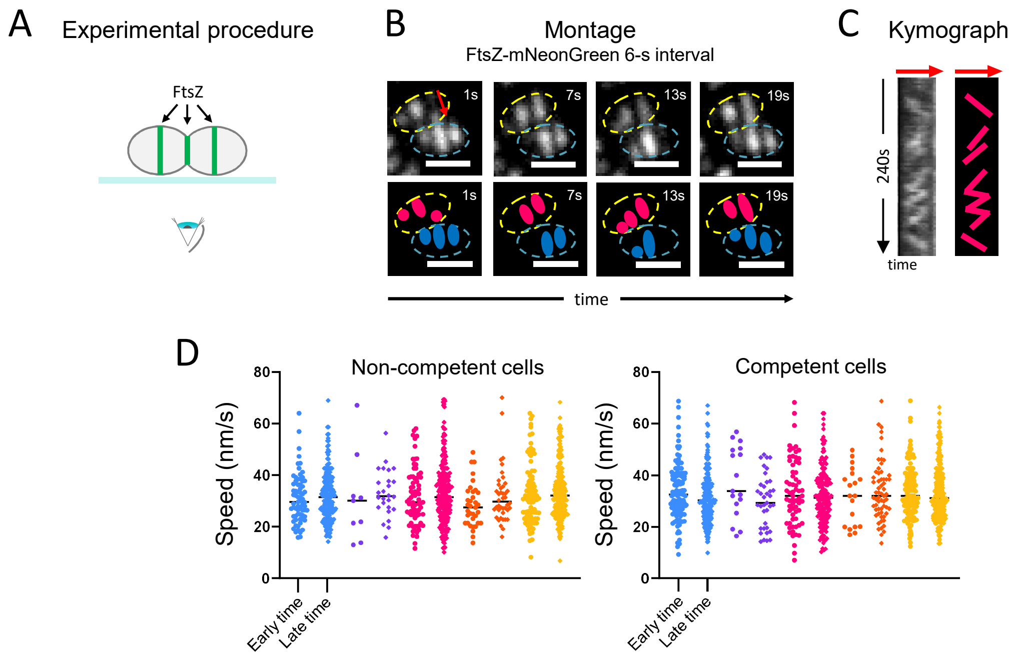


**Extended data Fig. 6: Dynamic of FtsZ remains similar at all stages of the cell cycle during competence.** Representative data are shown from two or three independent biological replicates.

**(A)** Schematics of the method to observe FtsZ-mNeonGreen in *S. pneumoniae* cells lying horizontally with the division plane orthogonal to the coverslip (related to Fig. 2A and Extended Data Fig. 3).

**(B)** *Upper part*: Montage of TIRFm images at 6s intervals showing FtsZ-mNeongreen. The red arrow indicates the trajectory extracted for kymograph analysis (voluntarily slightly shifted to the right to allow the visualization of FtsZ patches). *Lower part*: Cartoon representation of two FtsZ patches (red and blue dots). The contours of two cells are represented in yellow and turquoise dotted lines. Scale bars, 1 µm.

**(C)** *Left*: Kymograph from 1 to 240s obtained from the trajectory shown in B. *Right*: Cartoon representation of the kymograph.

**(D)** Speed of FtsZ-mNeonGreen recorded in different Z-ring type in non-competent (no CSP treatment) and competent cells at different time after competence induction (early time: 10-25 min after CSP addition; and late time: 30-55 min after CSP addition). A total of 1021 and 1038 trajectories were analyzed in non-competent cells and competent cells, respectively. One-way ANOVA statistical analysis indicate no significant differences between conditions.

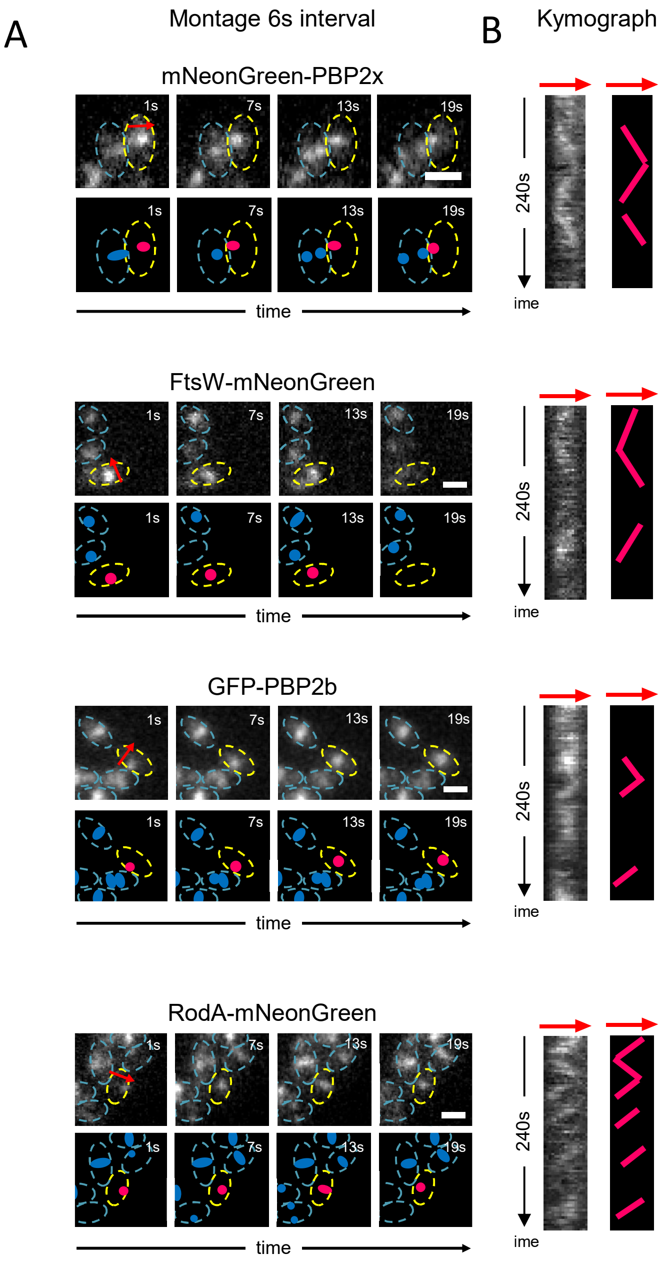


**Extended Data Fig. 7: Kymograph analyzes indicate that the speed of PBP2x and FtsW is reduced during competence compared to the speed of PBP2b and RodA.**

Representative data are shown from two independent biological replicates.

**(A)** *Upper parts*: Montage of TIRFm images at 6s intervals showing fusions of mNeonGreen or GFP with PBP2x (strain R4743), FtsW (strain R4728), PBP2b (strain WT gfp-pbp2b) and RodA (strain R4867). Red arrows indicate the trajectory extracted for kymograph analyses (voluntarily slightly shifted to allow the visualization of the fluorescent patches). *Lower part*: Cartoon representation of the fluorescent images (red and blue dots represent distinct fluorescent patches ). The cell contour is represented in yellow and blue dotted lines. Scale bars, 1 µm.

**(B)** Kymographs from 1 to 240s obtained from the trajectories shown in panel *A* (*left*), and cartoon representations (*right*). The slope of the fluorescent patches represented as red dots in panel *A* are shown.

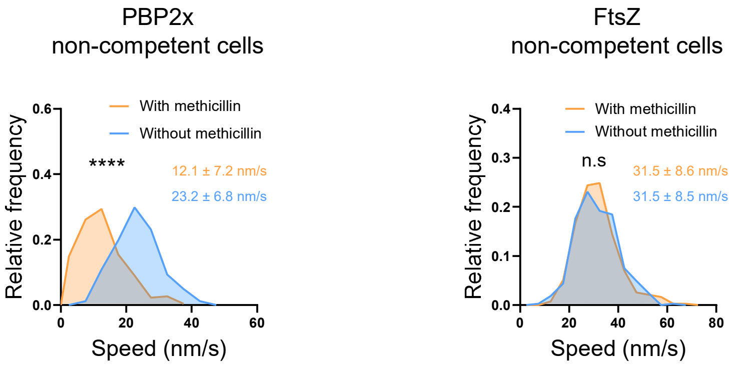


**Extended Data Fig. 8: Methicillin impacts the dynamics of PBP2x.**
Distribution of speed of mNeonGreen-PBP2x patches in a population of non-competent cells treated (310 trajectories), or not (268 trajectories), with methicillin and Distribution of speed of FtsZ-mNeonGreen patches in non-competent cells treated (430 trajectories), or not (385 trajectories), with methicillin. The average speed for each condition is indicated. Pairwise comparisons were done with a nonparametric Mann-Whitney test (****, P < 0.0001). Representative data are shown from two or three independent biological replicates.

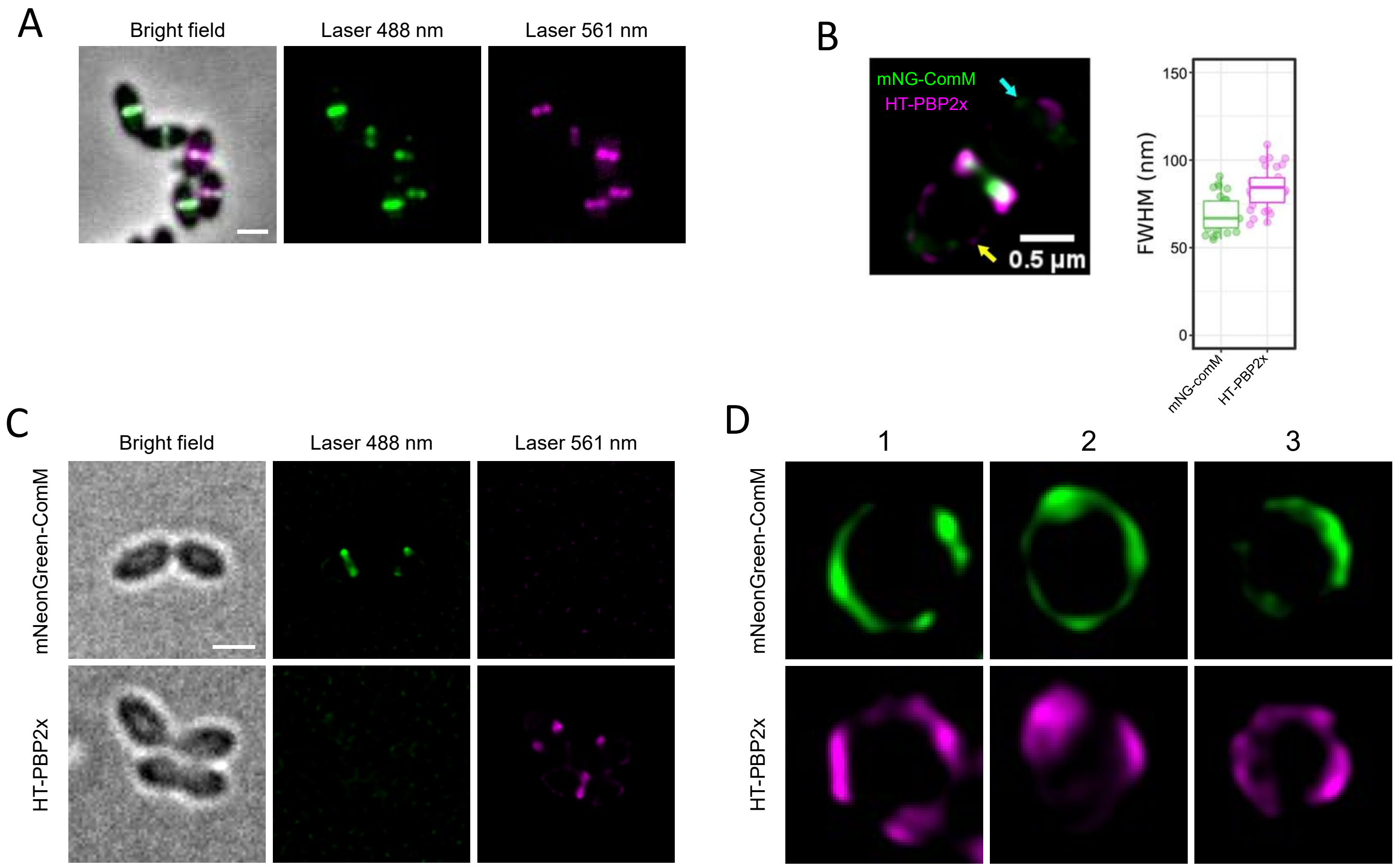


**Extended Data Fig. 9: Control experiments for co-localization analyses**

**(A)** Epifluorescence microscopy of the multicolor *mNeonGreen-comM, HT-pbp2x* strain R4746. Representative fluorescence and merged images between bright field (gray), mNeonGreen (Laser 488 nm, green), and Janelia fluor HaloTag ligand (Laser 561 nm, magenta) signals are shown. Superimposition of the ComM (green) and PBP2x (magenta) fluorescence signals appear false-colored in white. Scale bar is 1 µm.

**(B)** Estimation of lateral resolution in SIM² images. The full width at half maximum (FWHM) was calculated to determine the optical resolution in the lateral direction of dual color SIM² images. Data from strains R4746 were analyzed by fitting a Gaussian model function to the intensity profile in SIM² images. The average FWHM of mNeonGreen-ComM and HT-PBP2x were determined from small patches (cyan and yellow arrows for mNeonGreen-ComM and HT-PBP2x respectively) visible towards the cell poles (n=30 per condition).

**(C)** SIM² imaging of representative cells in bright field and two distinct channels (Laser 488 nm and Laser 561 nm).

**(D)** Individual channels of 3 representative rings analyzed in Sim².

Scale bar is 1µm.

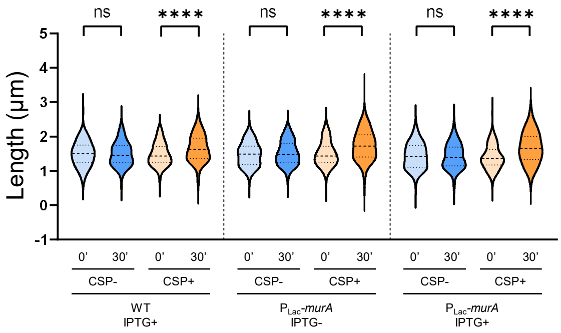


**Extended Data Fig. 10: Overproduction of MurA1 does not rescue the ComM-dependent cell elongation phenotype of competent cells.**

Violin plots representing the cell length distribution of non competent (CSP-) and competent cells (CSP+), measured on phase contrast microscopy images using > 100 cells for each strain. Cells were incubated with CSP for 30 minutes before imaging. 25th and 75th percentile are shown, with the horizontal line at the median.
